# Particle-based hydrogel inks and support matrices for biofabricating structural complexity, soluble gradients, and cell-lined channels in fully granular bioprinted systems

**DOI:** 10.1101/2025.06.07.658428

**Authors:** Julia Tumbic, Emily Ferrarese, Remington Martinez, Daniel Delgado, Thomas Ackleson, Christopher B. Highley

## Abstract

Towards achieving biomimetic complexity in biofabricated systems, an all-granular bioprinting system might use particle-based hydrogel inks to establish structures within a particle-based support matrix. In such a system, the granular support matrix can be designed to persist in the final construct and include cells incorporated prior to printing. To biofabricate complexity, bioprinting can introduce high-resolution heterogeneous structures that guide cell behaviors. The designs of the granular ink and support hydrogels are crucial to achieving complexity. High resolution structures and channels depend on small particles that flow and can be stabilized, and that can be printed and then removed, respectively. Herein, an all-granular system is described that used a granular formulation of an established, tunable hyaluronic acid-based hydrogel as the basis for a support matrix and a small particle gelatin hydrogel as an ink. Towards facilitating stabilization of the printed structure and flow during printing, the support and ink materials included soluble, interstitial components, and all exhibited yield stress behaviors characteristic of granular hydrogel systems. The support matrix’s viscoelastic properties were dependent on intraparticle hydrogel network design, and it could be stabilized against flow by photoinitiated crosslinking. The gelatin ink could form fine filaments, as small as 100 µm in testing here, and melted to leave channels within crosslinked support matrices. Channels could support flows introduced by hydrostatic pressure and could be used to rapidly transport soluble factors into the construct, which could be used to establish soluble gradients by diffusion and support cell viability. The all-granular system supported printing of complex, multimaterial structures, with feature resolution on the order of 100 µm and spatial positioning on the order of 10s µm. The process and materials exhibited biocompatibility with respect to cells included within the support matrix during printing or introduced into channels to begin establishing endothelialized bioprinted vessels.

## 1. Introduction

To mimic complex human tissues, with intricate physical and cellular architectures tied to tissue functions, tissue engineering approaches have turned to 3D bioprinting techniques. The development of this biofabrication technology has enabled multiscale control over construct complexity, from macroscale structure to microscale composition, and materials used in bioprinting approaches have spanned a range of formulations and applications(1–3). Bioprinting has been used to engineer tissues including bone (4–7), muscle(8–10), intestinal(11–16), and neural tissues(10,17). In addition to multiscale architectural control, biofabrication technology has been used to help build microphysiological systems(18,19). Research at the interface of biofabricating tissue systems and developing new biomaterial technologies is important to advancing capabilities for engineering biological structures.

The emergence of printing into permissive materials or support baths(20,21), including methods called embedded 3D printing or matrix assisted bioprinting, has addressed challenges in traditional extrusion-based bioprinting not easily resolved using other approaches(22–24). Printing an ink into a support matrix can allow complex structures like voids and overhangs to be fabricated and persist in the structure even after removal from the support post-printing(22,23,25). Alternatively, the support matrix can be designed as part of the final construct, serving as a 3D canvas into which cells and inks can be precisely positioned during the printing process(10,18,24,26–37). Designing the support matrix to participate in the final construct also enables the formation of features like discrete droplets(27), or voxels(28), of ink as well as perfusable channels(10,24,29–32,38) within the support matrix’s volume, creating a complex 3D system. Inks that may not have sufficient material properties to withstand gravity and have poor shape fidelity if printed traditionally can be successfully printed in conjunction with a granular support system(22,23). This technique expands the materials available for bioprinting by eliminating certain requirements for mechanical properties of the printing ink(39), allowing the use of softer materials where needed.

Support matrices vary in material choice, from synthetic polymers like photocurable Pluronic(24) to natural materials like modified hyaluronic acid(30) and composite ECM-based materials(31). They generally have two crucial rheological properties: 1) shear-thinning behavior, and 2) self-healing behavior. As an nozzle extruding ink moves through the support, the support must flow around the nozzle and to accommodate the ink, but re-solidify to encapsulate the ink. To achieve these properties, chemical bonding within the support material is often targeted for design. For example, hydrogels that are not covalently crosslinked by stabilized by supramolecular bonds can create a permissive environment, where bonds are readily and easily broken and reformed(27,30), thus satisfying both support requirements.

Granular hydrogels have been employed as promising support materials in matrix assisted bioprinting processes(22,23,26,40,41). A macroscale granular hydrogel is formed through packing or jamming of hydrogel-based microparticles, and these particle-based systems inherently have rheological properties crucial for matrix assisted bioprinting. In their jammed state, which occurs when particles are in contact with other particles(42,43), granular hydrogels behave like an elastic solid when at a standstill due to frictional forces between contacting particles(44). A force beyond the yield stress is required to overcome jamming within the particle network and fluidize the support(44,45). Upon cessation of this force, the particle network reforms and the support returns to a solid-like state(22,43). These repeatable and rapid solid-liquid and liquid-solid transitions are advantageous in their use as printable inks(46) as well as supports. Furthermore, any hydrogel that can be formed into microgels through methods like batch emulsification could be used as a granular support or ink, offering a generalizable approach to designing printable biomaterials.

Here, we demonstrate matrix assisted bioprinting using granular hydrogel materials as both the support and ink materials. The resulting constructs are granular hydrogels with printed features that can include perfusable vessel-like structures enabled by a removable granular gelatin ink. In applications as removable supports, granular materials have enabled printing retrievable objects within a particle-based support(22,23). Thay can also support perfusable channels in in examples that have included both granular hydrogels(29,31) and cell-based organoids(10) as particle-based support materials. Granular materials can also be used to support the printing of structures with high resolution features and specific cellular architectures (22,26), with particle-size influencing to printed feature resolution(10,47). As extrudable inks, granular hydrogels have been shown to enabling printing of biomaterials that are not easily printed if they are not formulated as granular materials(46,48). Extruded granular materials are often stabilized by physical forces after deposition, but can be further stabilized through a secondary interparticle crosslinking reactions(49–51) or through inclusion of a crosslinkable interstitial material(10,29,52). As with granular supports, resolution of printed structures using granular inks will be influenced by particle size. In employing granular materials as supports and inks, work described below develops a removable granular ink.

By printing granular inks into a granular support, we aimed to establish a highly tunable 3D biomaterial system, or canvas, that would support biofabrication specific and customizable architectures for uses in tissue engineering. Granular hydrogel formulations allow biomaterials that are well-established for engineering biomimetic complex to be printed, would allow high-resolution features to be approached using microgels with small diameters(47) – here, in the low 10s of microns range – and would allow deposited inks to serve as supports themselves after deposition. The readdressing of printed locations would allow complex features to be printed.

Two commonly used biopolymers were used as the basis for our granular hydrogel support and inks: hyaluronic acid and gelatin. Hyaluronic acid (HA), found in tissues like the brain and articular cartilage, modified with norbornene functionalities (NorHA) is established as a robust biomaterial for forming complex hydrogels(53). NorHA can be crosslinked and modified through well-controlled photoinitiated thiol-ene chemistries (53,54). As particle-based support material, NorHA served here as the 3D biomaterial canvas. Gelatin is widely known for its thermal gelation and has been used as a continuous ink(10,55) as well as in granular formulations in biofabrication(23,38,41,56) – here we used it as a granular ink that could be removed, if desired, by heating. Both materials were made into microgels and then jammed as granular hydrogels. As the formulations of the NorHA and the gelatin hydrogels as granular hydrogels consisting of small particles and containing interstitial polymers are new, we first characterized their mechanical and dynamic properties through rheology. We then show that granular inks can be extruded into a granular support to facilitate the formation of perfusable channels, high resolution depots (or voxel) with controllable spacings, branching structures, and depots surrounding a filament. Finally, we illustrate that cell viability within the support material can be maintained or enhanced in a thick scaffold with channels, compared to a construct without channels, and that endothelial cells can be added to a channel as a step towards vascularizing all-granular hydrogel systems.

## 2. Methods

### 2.1 Synthesis

NorHA was synthesized using a previously described method(53). Briefly, to enable modification of HA with norbornene in an anhydrous reaction, sodium hyaluronate (NaHA, 60k, Lifecore) was converted to acid form by dissolving NaHA in deionized (DI) water to achieve a 2 wt% solution, followed by addition of an ion exchange resin (Dowex, Sigma). This material was then filtered over a Buchner funnel to remove resin, and the pH of the HA was titrated to 7 using tetrabutylammonium hydroxide (TBA-OH) prior to lyophilizing the solution. The resulting TBA-modified HA, was then dissolved in anhydrous dimethylsulfoxide (DMSO, Sigma) and reacted with 5-norbornene-2-carboxylic acid and 4-(dimethylamino)pyridine (DMAP, Sigma), catalyzed with di-tert-butyl decarbonate (Boc2O, Sigma), under nitrogen. The reaction was run overnight. The resulting material was dialyzed against DI water for three days, followed by purification via ethanol precipitation, resuspension in DI water, and further dialysis against DI water for an additional three days. This material was lyophilized and stored at -20°C until needed. The degree of HA modification with norbornene as a fraction of total disaccharide units was measured via H^1^ NMR in D2O (SI Fig. 1).

**Figure 1:**
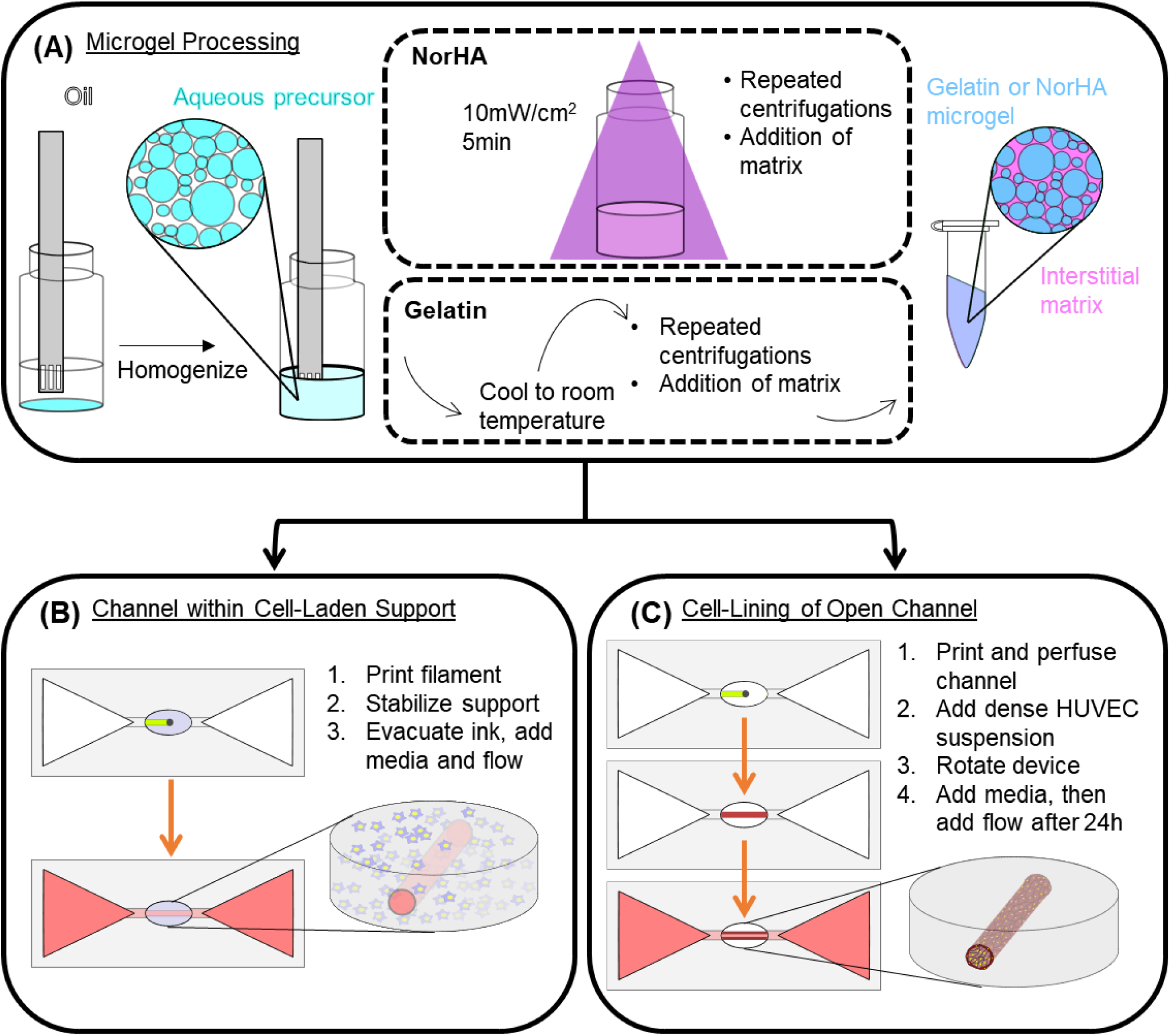
Schematics of the (A) NorHA and gelatin microgel processing, (B) the experimental process for creating a perfusable channel within a fibroblast-laden NorHA support, and (C) the experimental processing to creating a HUVEC-lined open channel.

### 2.2 Support medium and granular ink preparation

A precursor 3 wt% NorHA solution was prepared in phosphate-buffered saline (PBS) containing approximately 1mM, 2mM, or 4mM dithiothreitol (DTT, Sigma) and 6.6mM lithium phenyl-2,4,6-trimethylbenzoylphosphinate (LAP, Sigma). Control of the DTT concentrations lead to crosslinking densities of 2 mM, 4 mM, and 8 mM of norbornene groups consumed in the crosslinking reaction, and the materials are referred to as NorHA-2, NorHA-4, and NorHA-8, respectively. Precursor NorHA solutions were homogenized in light mineral oil with 2% Span 80 for 2 minutes at 3000 rpm, followed by UV curing for 5 min at approximately 10 mW/cm^2^ (Fig. 1A). The microgels and oil were then stirred for 2-5 min using a stirplate at 1000 rpm with excess isopropanol to begin de-emulsification. After stirring, the mixture was filtered over a 0.22 µm hydrophilic PVDF membrane. Microgels were washed an additional two times over the filter with isopropanol. After filtering, the microgels were left to dry on the benchtop under ambient conditions for roughly 5-10 minutes then added into a 50 mL conical and vortexed with 70% ethanol and placed on a rocker overnight. To finish processing, microgels were washed with PBS four times using centrifugation to ensure ethanol removal and equilibration in PBS. Microgels were then filtered over a 40 µm cell strainer and stored at 4°C until needed.

To prepare the NorHA support, the NorHA microgels were centrifuged at 3214 rcf for 10 minutes, and PBS was decanted. A small volume of PBS was then added back to loosen the microgel pellet, and the microgels were then aliquoted into Eppendorf tubes, and centrifuged at 5283 rcf for 5 minutes. Again, excess PBS was removed. An interstitial matrix solution was then added to these microgels. To prepare this solution, a 1 wt% NorHA solution was prepared in PBS with 0.625mg/mL of an MMP-degradable dithiol crosslinker (GCNSVPMSMRGGSNCG), synthesized using a peptide synthesizer (Liberty Blue), and 6.6 mM LAP. This interstitial matrix was added in equal volume to the NorHA microgels in the Eppendorf tube. The microgels and interstitial solution were vortexed for 20-30 minutes, then centrifuged at 5283 rcf for 5 minutes. Excess 1% NorHA solution was then removed, and centrifugation was repeated if necessary to remove excess solution. The final material was designed to be a jammed granular NorHA hydrogel with spaces between the particles including the interstitial material to support interparticle crosslinking in the final construct.

Gelatin microgels were prepared using a 15 wt% gelatin solution in PBS and a light mineral oil solution with 2% Span 80 that were separately heated to 80°C . The oil was then added to the gelatin solution and homogenized at 9000 rpm for 2 minutes. The mixture was then stirred using a stirplate at 1000 rpm until the emulsion reached room temperature. To prepare the gelatin-based ink, the microgels were jammed by centrifugation at 5724 rcf for 2 minutes. A 1% HA solution was prepared in PBS and then added to the gelatin microgels in a volume equal to that of the gelatin microgels and then mixed together. All washing steps and storage conditions were the same as for the NorHA microgels.

### 2.3 3D printing device preparation

Devices to hold the granular hydrogel during printing and perfuse medium into printed channels were created from polydimethylsiloxane (PDMS, Sylgard-184). The PDMS devices were formed by casting them off custom-designed molds and frames that were printed using a Form 2 printer (Formlabs). The device design consisted of three reservoirs in series connected by channels. After printing, the molds were processed by soaking twice in isopropanol followed by post-curing at an elevated temperature for at least an hour. The mold, which defined the wells, was glued onto a glass slide, and a frame, which defined the outer edges of the PDMS device, was secured onto the glass slide using binder clips. PDMS was prepared by mixing a 1:10 ratio of curing agent to base and then cast into the frame and around the mold and cured at 37°C overnight. After removal from the molds, PDMS devices were bonded to glass slides via plasma treatment in air. Shortly after, these devices were placed into a solution of 1:200 v/v 3-mercaptopropyltrimethoxy silane to ethanol for approximately 1 hour, then washed with ethanol and placed into 1M HCl for at least one hour prior to washing with DI water(57).

### 2.4 Particle size analysis

NorHA microgels were tagged with a thiolated rhodamine-B (Rho-B) peptide via photoinitiated crosslinking to of the free thiol on the peptide to pendant norbornene groups on the microgels that were not consumed during crosslinking. To fluorescently image gelatin microgels, FITC-dextran (2 MDa, Sigma) was added to the gelatin solution prior to homogenization. A Leica DMI-8 microscope was used to image the microgels using a 20x objective. Microgels were diluted with PBS and placed onto a glass slide. A coverslip was then placed over the microgels. The fluorescent images were thresholded in ImageJ, then the watershed function was used to separate contacting particles, followed by using the particle analysis function to obtain microgel diameters. Particles that were unable to be separated using the watershed function were discarded prior to running particle analysis. Images of the undiluted jammed support were taken as well images of the jammed material on a glass slide beneath a glass coverslip.

### 2.5 Rheology

Flow, frequency, and strain sweeps were carried out on both the NorHA support medium and gelatin microgel ink. A parallel plate system was used, as this provides a gap 10x the microgel diameters(44). To prevent wall slip, a sandblasted 20mm plate with a solvent trap was used as the top geometry, and sandpaper was placed on the bottom plate. Gap size was set to 500µm, and temperature was kept at 20°C. Prior to each test, the NorHA and gelatin granular hydrogels were sheared for 150 s^-1^ and 250 s^-1^, respectively, to erase shear history or aging and reset the material. For flow sweeps, the shear rate was decreased from 500 s^-1^ to 0.001 s^-1^ with steady-state sensing with 5% tolerance over a 5 s sampling period and a max equilibration time of 60 s. The stress to strain rate relationship was fit to the Herschel-Bulkley equation, which allows modeling the materials as shear-thinning fluids – when flowing – with a yield stress:

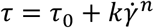

where 𝜏 is shear stress, 𝜏_0_ is the yield stress, k called the consistency index, 𝛾̇ is shear rate, and n is a constant describing flow behavior. When n < 1, the material is shear thinning under flow conditions, for n > 1, the material is shear thickening, and when n = 1 the material is a Bingham plastic. Frequency sweeps were carried out from 0.01 to 10 Hz with an applied strain of 1%, and strain sweeps were done from 0.01% to 500% with an applied frequency of 1 Hz. For the NorHA support, a photocuring step was included using a quartz bottom plate that allow light filtered to 320-390 nm to be applied to samples. The test was conducted at room temperature with an applied strain and frequency of 1% and 1 Hz. A delay time of 2 minutes was used to obtain pre-photo exposure storage and loss moduli. The material was exposed to light with an intensity of 10 mW/cm^2^ for 60 s. After photo exposure, the time sweep was run for an additional ten minutes at 1 Hz application of 1% oscillatory strain.

### 2.6 3D printing

To create perfusable channels via 3D printing, the PDMS devices described earlier were used. The PDMS channels connecting the side reservoirs to the central reservoir were blocked with 15% gelatin to isolate the NorHA support to the central reservoir. The NorHA support was then added to the central reservoir. A gelatin ink was prepared and loaded into a 100 µL Nanofil syringe. This syringe was loaded onto either a Felix 3.2 3D printer modified for syringe extrusion(58), or a Felix Bioprinter with custom-made syringe adaptors designed in AutoCAD and printed with a Form 2 printer (Formlabs).

Ink extrusion was controlled in G-codes by modifying the filament extrusion parameter E while maintaining a feedrate, which corresponds to the nozzle movement rate, of 50 mm/min. A 26G needle was used for all prints except for the lowest flow rate tested, where a 36G needle was employed. The NorHA support was then photocrosslinked at 10 mW/cm^2^ for 1 minute. In instances where gelatin ink was removed, the device was placed in a humidified environment at 37°C, with one side reservoir containing PBS to create hydrostatic pressure at one end of the channel that facilitated removal of the gelatin after melting. Images were taken both after printing and after ink removal on an inverted widefield microscope (DMi8, Leica) with a 5x objective. Measurements of filament diameter, channel diameter, and perfusable diameter were made using ImageJ. Statistical analysis was done in OriginPro 8.1, with a one-way ANOVA and post-hoc Tukey test to determine statistical significance.

To print depots and other complex features, NorHA support was added to a PDMS holder, and G-codes were created that corresponded to the target design. Gelatin inks including either dilute fluorescein isothiocyanate (FITC)-labeled dextran or rhodamine isothiocyanate (RITC)-dextran within the gelatin microgels were used for imaging purposes. The inks were printed either alone or in multilink processes. In these prints, where removal of the gelatin structures was not desired, the printed constructs were maintained at room temperature, below 37°C. After printing, images were taken on with a 5x objective.

### 2.7 Diffusion

After printing channels and removing the solubilized gelatin, PBS in the reservoirs was replaced and the device equilibrated for 1 h at 37°C prior to initiating the diffusion experiments. A 1 µg/mL Rho-B (Sigma) solution and a 0.112 mg/mL FITC-labeled bovine serum albumin (FITC-BSA, Sigma) were prepared in PBS. The corresponding solutions were added in equal volumes to both side reservoirs, and either the microscope began continuous image acquisition for the Rho-B experiment, or timepoints of 1 h, 3 h, 6 h, and 24 h were taken for the FITC-BSA experiment. Both experiments were carried out at room temperature. The fluorescence of the hydrogel systems, which changed as the fluorescent molecules diffused out of the channel into the support material, was acquired in the images using controlled exposure time and light intensity that were quantified using ImageJ.

### 2.8 Material preparation for cell culture experiments

The granular NorHA support material and gelatin ink were prepared in the same manner as described earlier, but in a sterile environment in a laminar flow hood after microgel fabrication and washing in 70% ethanol. The 1% NorHA matrix was made with complete fibroblast media and then sterile-filtered through a 0.22 µm PVDF syringe filter. A thiolated arginylglycylaspartic acid peptide (GCGYCRGDSPG, GenScript) was added to the 1% NorHA to a final concentration of 1 mg/mL. The 1% HA for the gelatin ink was exposed to germicidal UV for sterilization. PDMS devices were sterilized by soaking in 70% ethanol after treatment with silane was complete.

### 2.9 Preparing thin support gels containing cells and measuring proliferation

Fibroblasts (mouse, NIH 3T3s) were cultured, with plating densities of 2000 cells/cm^2^ used in passaging. Dulbecco’s modified eagle medium (DMEM) containing L-glutamine and high glucose was supplemented with 10% calf bovine serum and 1% penicillin/streptomycin. Cells were harvested using 0.05% trypsin with EDTA and centrifuged at 130 rcf for 5 minutes. For experiments in granular hydrogels, cells were resuspended at a concentration of 100 million cells/mL after centrifugation.

To assess cell viability within the NorHA support, cell suspensions (100 million cells/mL) were added to the NorHA-4 support at volumetric ratios of 1:10, 1:20, and 1:50 cell suspension:NorHA support. 100 µL of this cell-containing support was added to wells in a 48 well plate and then photocrosslinked at 10mW/cm^2^ for 60 s. Complete media was then added to each well. Proliferation was quantified using the Alamar blue assay on days 1, 3, 5, 7, and 9, with fold change calculated relative to day 1. The assay was prepared according to manufacturer’s instructions, where the assay’s reagent solution was mixed with complete medium in a 1:10 volumetric ratio, heated to 37°C prior to addition to hydrogel samples.

### 2.10 Preparing thick support gels with channels for studying effects of channels on cell viability

Cells were harvested and added to NorHA-4 support as described above. The PDMS device used for introducing medium into printed channels was prepared as described above. The cell-laden support was then added to the central reservoir of the printing device, shown in Fig. 1B. To create constructs without channels, the NorHA support was photocrosslinked, as described earlier, and complete media was added to the side reservoirs and on top of the support.

Channels in constructs were prepared by printing a filament of granular gelatin ink into the cell-laden support, photocrosslinking the support, and then removing the gelatin ink by incubating the device in an incubator at 37°C.

Cell viability was tested by removing the constructs from the device and cutting the granular hydrogel perpendicular to the channel expose the cross section, then staining with calcein blue AM (Invitrogen) and ethidium-homodimer 1 (EthD-1, Invitrogen). Prior to staining, the cross sectioned constructs were washed with Dulbecco’s modified PBS (DPBS), then viability stains were introduced in a DPBS solution containing 20 µM calcein blue AM and 4 µM EthD-1 was added to the samples and placed in the incubator for at least 20 minutes. After incubation, samples were washed with DPBS then imaged in an inverted widefield microscope (DMi-8, Leica).

### 2.11 Lining of channel lumen with endothelial cells

Human umbilical vein endothelial cells (HUVECs) were cultured with the EGM-2 Bulletkit (Lonza). Passages were performed with 0.025% trypsin/0.01% EDTA in PBS, followed by centrifugation at 200 rcf for 5 minutes and counted using a 1:5 dilution with trypan blue. For experimentation, HUVECs were used at passage 5.

To seed onto channel walls, a perfusable channel was first printed into an acellular NorHA-4 support as described earlier. The interstitial 1 wt% NorHA was prepared with thiolated RGD (GCGYCRGDSPG, GenScript) at a final concentration of 8mg/mL. After removing the gelatin ink used in the channel printing process, HUVECs were harvested and resuspended at a concentration of 30 million cells/mL. Following a previously published protocol(59), 5-10 µL of the cell suspension was added to the channels, then the devices were placed in the incubator and were flipped every 30 minutes for 4 hours (Fig. 1C). Media was then carefully added to the reservoirs. After approximately 24 h, flow was to the cell-lined channels by placing the device on an orbital rocker.

Fluorescent imaging was done by staining the HUVECs with CellTracker Deep Red (Invitrogen) on days 1 and 3 post-seeding. The channels were washed with DPBS, then a roughly 5 µM CellTracker Deep Red stain solution was added to each sample and the devices were incubated for 15 minutes at 37°C. This stain solution was then removed and replaced with DPBS. Channels were imaged on a Leica DMi8 using 5x and 10x objectives.

## 3. Results

### 3.1 Characterizing granular support and ink materials

Particles with average diameters on the order of 10 – 20 µm were created for both the granular ink and support materials from batch emulsification. NorHA particle sizes and size distributions were dependent on the crosslinking density of the microgels for the same processing parameters. Particle populations were polydisperse, as expected from batch emulsification. Average particle diameter increased and size distributions broadened with decreasing crosslinking (Fig. 2A-B and Table 1). NorHA-2 and NorHA-4 particles had similar particle diameters and size distributions (Fig. 2B and Table 1), with the diameter becoming less and size distribution narrowing for NorHA-8. Qualitatively, when jammed, the smaller NorHA-8 particles were less clear compared to jammed NorHA-4 and NorHA-2 particles (Fig. 2C), as would be expected from increased light scattering with smaller particle size.

**Figure 2:**
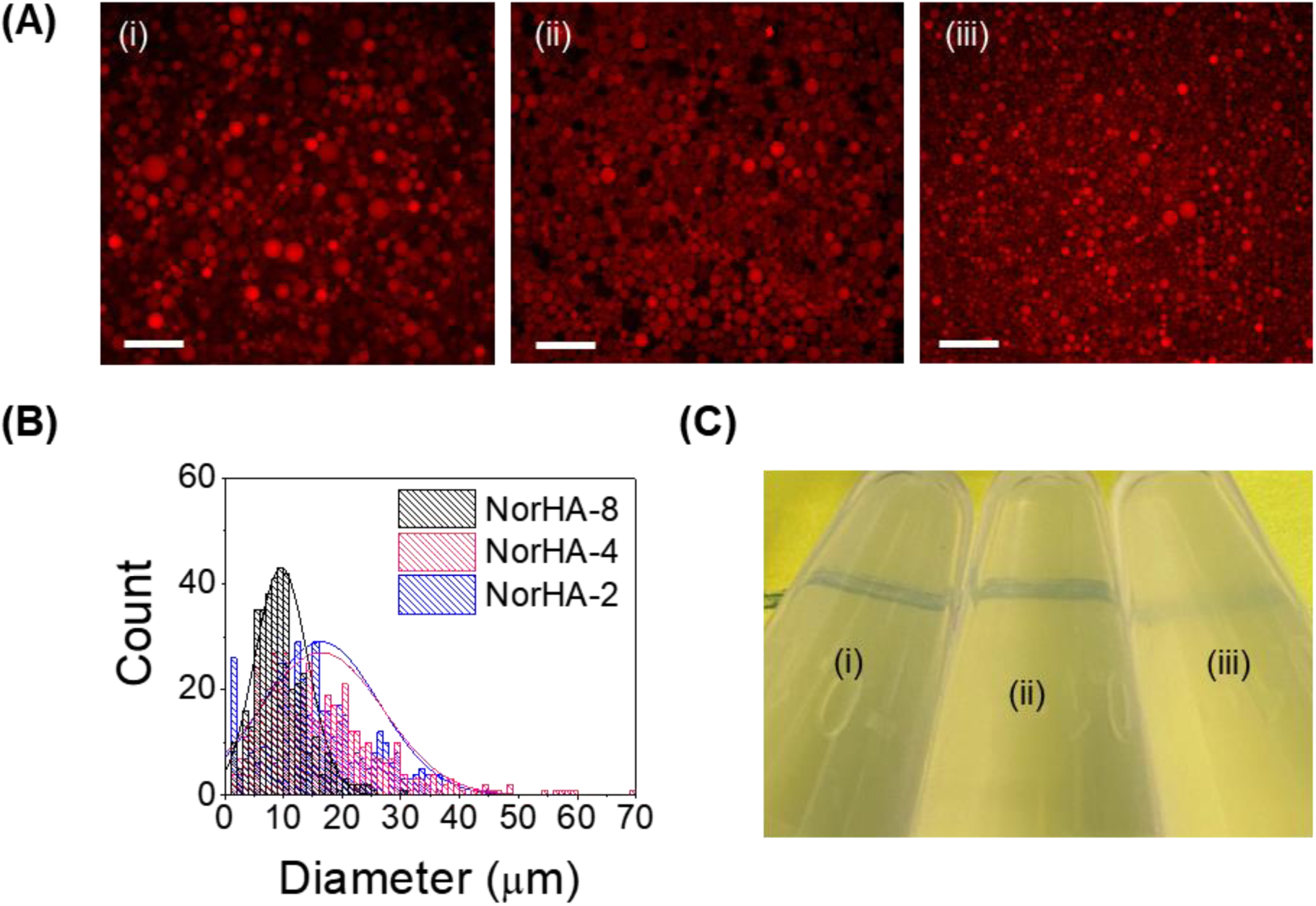
(A) Fluorescent images of NorHA particles of different crosslinking densities of (i) 2mM, (ii) 4mM, and (iii) 8mM, where red is the Rho-B peptide (scalebars: 100 µm); (B) particle size distributions of each NorHA microgel group; (C) images of jammed (i) NorHA-2, (ii) NorHA-4, and (iii) NorHA-8 particles in the bottoms of microcentrifuge tubes over a black line to illustrate relative transparencies.

Rheological measurements were made on jammed granular hydrogels comprised of NorHA-2, NorHA-4, and NorHA-8 particles (Fig. 3). Unidirectional shear data show yield stress behavior for all hydrogels (Fig. 3A), illustrated by the plateau in stress at low shear rates. This yield stress was dependent on the extent of NorHA crosslinking within microgels, with lower crosslinking leading to a linear decrease in yield stress. The Herschel-Bulkley equation was fit for all materials tested (SI Table 2).

**Figure 3:**
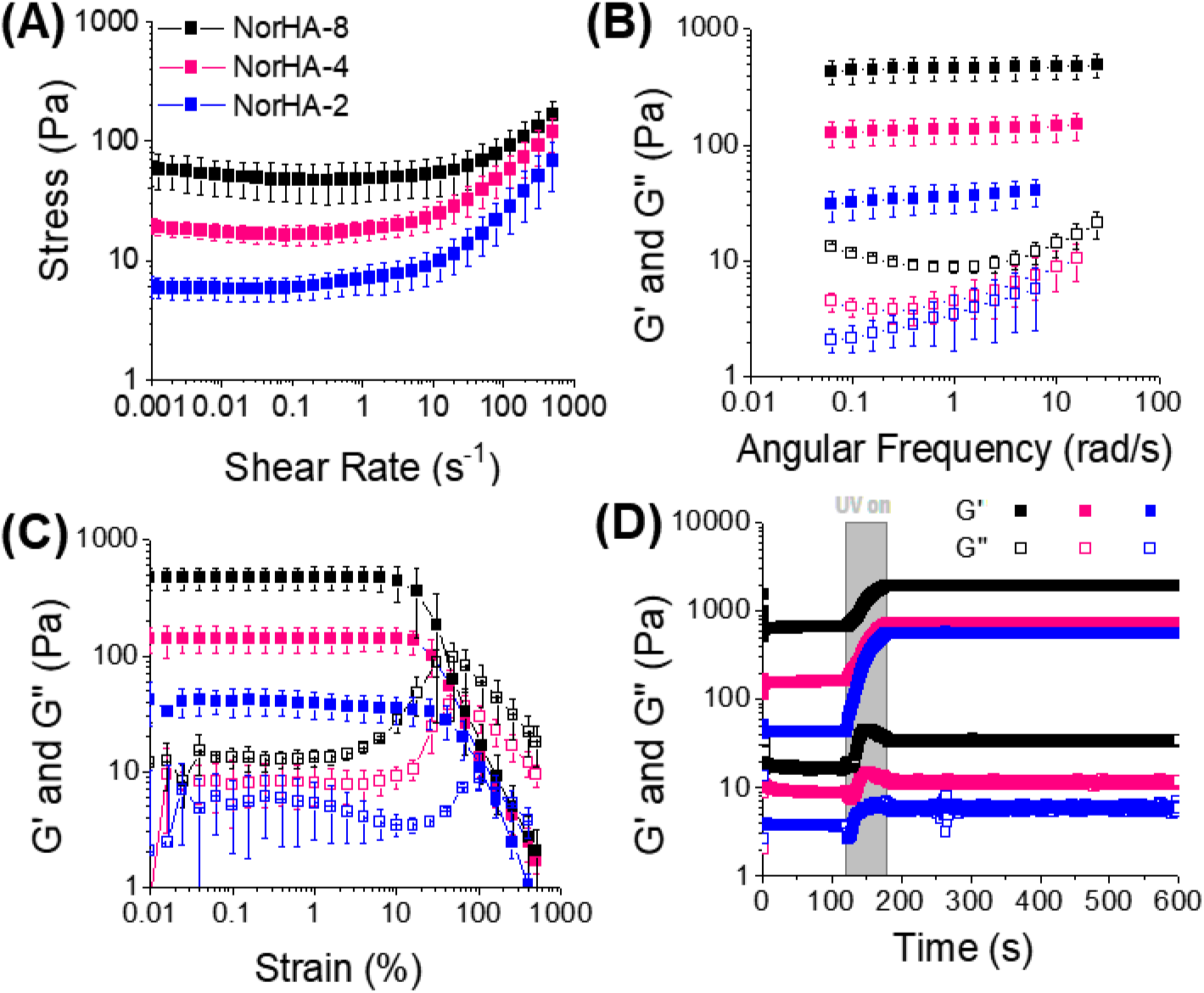
Rheological characterizations of jammed NorHA-2, -4, and -8 supports, with (A) flow sweeps, (B) frequency sweeps, (C) strain sweeps, and (D) UV curing tests conducted. Error bars represent standard deviation.

The NorHA systems exhibited behaviors typical of jammed granular hydrogels, as seen in frequency sweeps (Fig. 3B). Specifically, the storage moduli for the NorHA-4 and NorHA-8 systems were frequency-independent, while the NorHA-2 storage modulus showed a slight dependence on frequency. A characteristic minimum was present in the loss modulus frequency sweeps, which appeared at decreasing frequency with decreasing crosslinking density of the particles.

Prior to initiating interparticle crosslinking, strain sweeps showed that the storage moduli for all the NorHA systems were strain-independent at low strains (Fig. 3C). Under increasing strain, granular hydrogels yielded exhibited a transition to fluid-like behavior, the onset of which depended on the internal crosslinking density of the microgel particles. As the particle crosslinking increased, the strain at which yielding was observed increased. The G’-G” crossover followed this trend as well. After initiating interparticle crosslinking, stiffnesses of the granular hydrogel systems were dependent on crosslinking density of the microgels (Fig. 3D), as expected given bulk properties are dependent on the microgel properties in granular systems. MMP-degradable crosslinker concentrations included in the interstitial material were constant for all samples, so differences in properties observed are a function of the microgel populations that form the granular hydrogel systems. Final modulus values ranged from approximately 450 Pa for the NorHA-2 system to roughly 1.7 kPa for the NorHA-8 system. The moduli of the granular systems increased when interparticle crosslinking was initiated (SI Fig. 2).

The granular gelatin hydrogel had an average particle diameter of 4.6 ± 3.77 µm with a coefficient of variation of 82% (Fig. 4A and 4B, and Table 1), reflecting polydispersity resulting from the emulsification process used. The gelatin ink was shear-thinning and easily extruded through a 25G needle (4C). Similar to the NorHA microgel systems, the gelatin ink had a yield stress (SI Fig. 3), and exhibits a frequency-dependent loss modulus with a characteristic minimum (Fig. 4D). The granular gelatin ink yielded at strain of 24.55%, measured in oscillatory strain sweeps (Fig. 4E and Table 3). Below its melting point, at room temperature, this ink could hold a shape as a thick droplet, but upon heating to 40 °C, this droplet liquefied due to gelatin particles melting (Fig. 4F).

**Figure 4:**
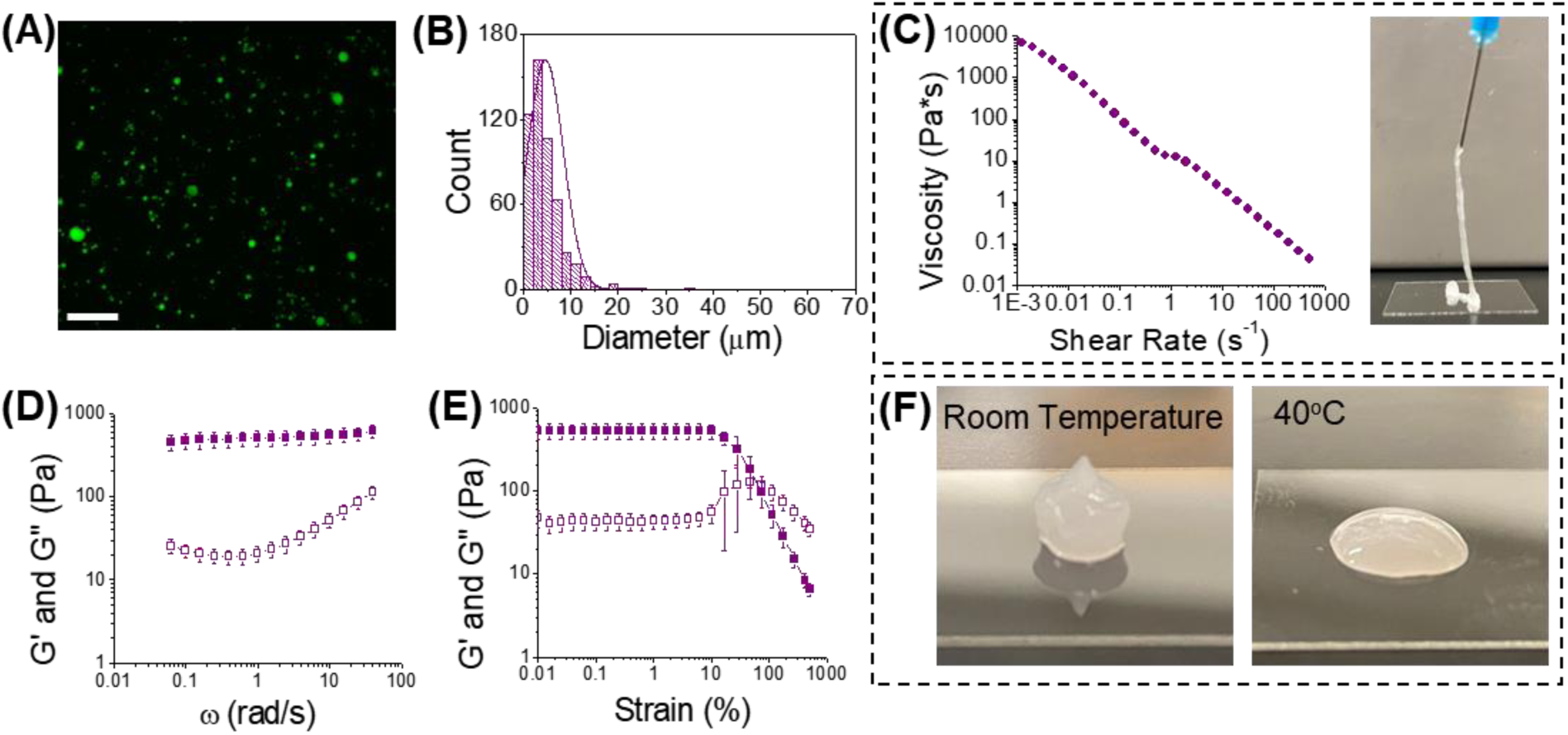
Characterization of gelatin microgels, with (A) fluorescent images of gelatin/FITC-dextran microgels taken (scalebar: 100 µm) and (B) the particle size distribution measured; rheological characterizations show (C) shear-thinning behavior, also shown by the ink flowing through a 25G needle, (D) frequency sweeps and (E) strain sweeps were conducted as well; qualitatively, (F) the ink can hold a thick droplet shape at room temperature, but upon heating to 40°C, the ink melts. Error bars denote standard deviation.

### 3.2 3D printing complex material architectures and channels in all-granular hydrogel systems

To determine what resolution of 3D printed features might be achieved within our all-granular hydrogel system, filaments were printed with varying flow rates to control filament diameter. Gelatin ink in which the particles contained high molecular weight FITC-dextran was used for imaging purposes. Widefield fluorescent images were taken directly after printing to visualize printed features. Filaments with diameters ranging from 100 µm up to 700 µm were printed (Fig. 5A). To print the smallest diameter filament, a 36G needle was used, the other filaments were printed with a 26G needle.

**Figure 5:**
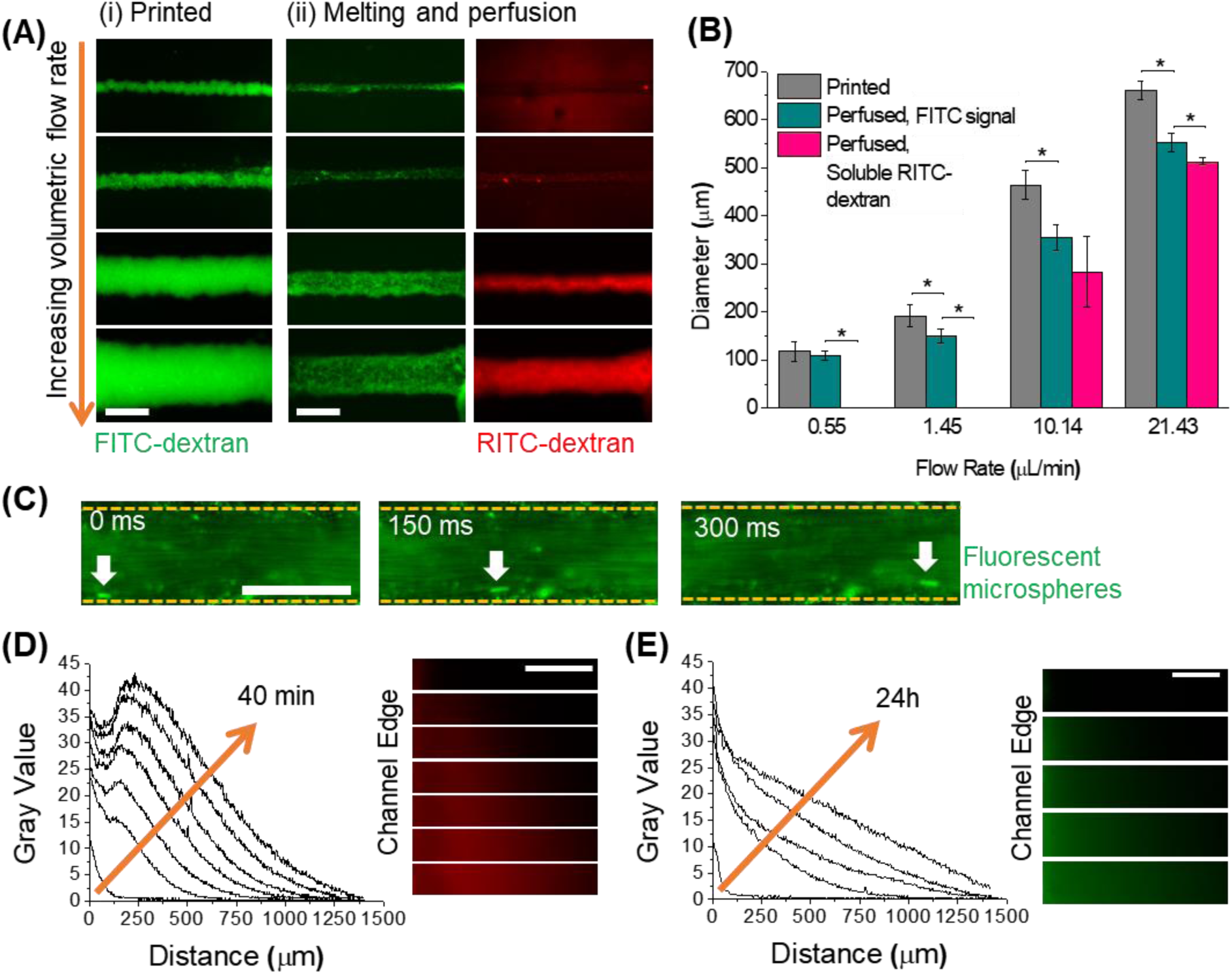
(A) Fluorescent images of (i) printed filaments and (ii) post-thermal treatment to 37°C to melt the ink, with green representing FITC-dextran and red representing RITC-dextran (scalebar: 400 µm), (B) quantification of printed filaments compared to post-thermal treatment of the channel with ink and the perfusable area only (* indicates a significance of p<0.05 and error bars denote standard deviation), (C) flow of fluorescent microspheres through a perfused channel, showing laminar flow (scalebar: 300 µm), (D) diffusion of a Rho-B solution from a channel into the surrounding gel overtime (scalebar: 300 µm), (E) diffusion of a FITC-BSA solution from a channel into the surrounding gel over time (scalebar: 300 µm).

To then establish channels, the gelatin was removed by incubation at 37 °C; constructs were imaged two days after the original print. High molecular weight FITC-dextran could be visualized on the channel periphery after crosslinking the granular support matrix and evacuating the melted gelatin from the channels – a result of construct crosslinking immobilizing a fraction of the gelatin particles via interstitial HA at the filament periphery prior to melting. The FITC-dextran signal allowed quantification of channel diameters after melting gelatin and incubating the construct over two days. The channel diameters, as measured from the FITC signals, were observed to significantly decrease in all groups except the smallest channel. During perfusion of the channels with media containing RITC-dextran, the channel diameters as measured by the RITC signals were statistically similar to those measured from the FITC signals (Fig. 5B). The two smallest channels, however, were not observed to support perfusion driven by hydrostatic pressure in the devices used.

Once channels were established, the ability of perfusable channels to support laminar flow was visualized using fluorescent microspheres (FluoSpheres, 1 µm, yellow-green, polystyrene, Molecular Probes). To induce flow through the channel, medium containing a suspension of fluorescent microspheres was added to one of the reservoirs with the other reservoir initially empty. Images captured over time (Fig. 5C) showed that the microspheres were convected in a laminar flow pattern and did not get trapped along the walls of the channel, despite microscale irregularities along the walls of the channel due to processing using granular materials.

Perfusable channels offer the potential to create diffusive gradients within a material. To test the ability of small molecules and large proteins to diffuse into the surrounding support from a channel, solutions of RhoB and FITC-BSA were added to the reservoirs of the devices and the fluorescence was monitored over time via microscopy. As expected, RhoB diffused quickly into the surrounding NorHA support, reaching distances of approximately 1400 µm from the channel edge over a span of 40 minutes (Fig. 5D). On the other hand, FITC-BSA predictably diffused much slower, with the FITC-BSA reaching approximately the same distance over a 24 h period (Fig. 5E).

To illustrate the versatility of this printing system and show the potential to mimic complex architectures found in biological systems in vitro, non-removable features were printed into complex patterns within the NorHA-4 support. Fluorescent polymers were separately added to the gelatin microgels during particle fabrication, then used to visualize the printing of two different inks. To demonstrate structures that would be difficult to fabricate by other means, a filament was first printed surrounded by discrete depots of a second ink (Fig. 6A). Branching structures that could be used to form more complex biological structures were also printed, with filaments of roughly 200 µm in diameter (Fig. 6B). Voxel-like depots of two different fluorescent inks, approximately 300µm in diameter, were extruded in a regular, alternating pattern (Fig. 6C). Additionally, placement of depots with varying proximities to one another was easily controlled, as show by printed depots with a spacing of approximately 600 µm, depots right next to each other, and depots printed within one another (Fig. 6D).

**Figure 6:**
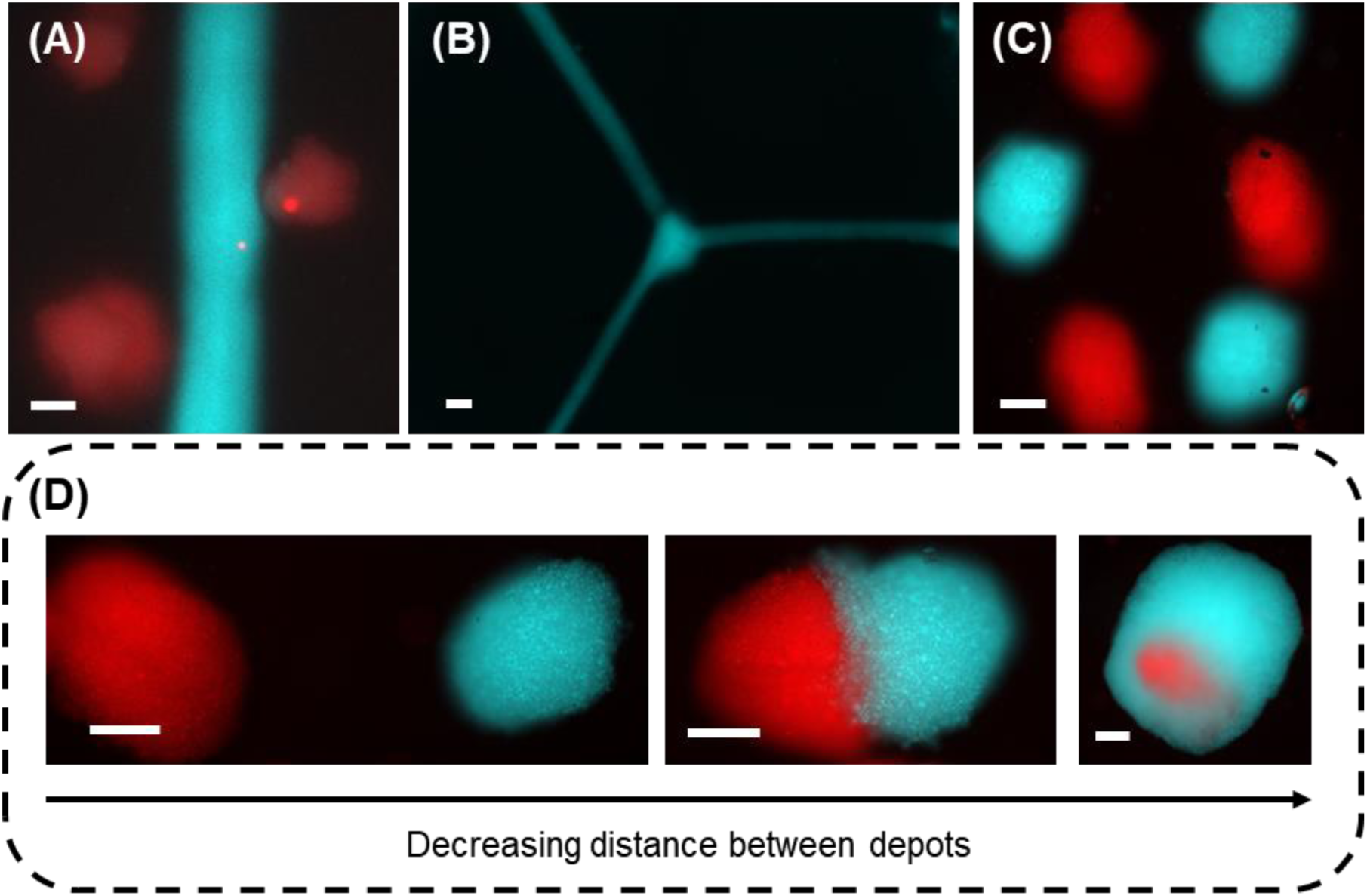
printing of complex features within a jammed NorHA microgel support, with (A) depots surrounding a filament, (B) a branching structure, (C) voxel-esque printing of depots, and (D) two depots printed at varying distances from one another. Scalebars: 300 µm.

### 3.3 Including cells within all-granular hydrogel bioprinted constructs

To observe cell growth within these materials, fibroblasts were seeded into NorHA-4 microgel support gels, as described. To ensure adequate nutrient and oxygen transport, constructs used were 100 µm thick and were cultured over a nine-day period with proliferation testing every other day (SI Fig. 4A). All samples showed either an increase or maintenance of metabolic activity over the testing period. The increases in metabolic activity (fold change values, SI Fig. 4B) were dependent on cell density. The lowest cell density (1:50 cell suspension:support matrix) had the larger increase in fold change, with the highest cell density (1:10) had the lowest fold change increase.

Having observed cytocompatibility in increasing metabolic activity in thin constructs, we next looked at whether channels would support viability in thick constructs. Cell viabilities in constructs with thicknesses of 3-5 mm with and without channels were compared with the expectation that channels would support higher viability compared to a solid construct. A dense suspension of fibroblasts was added to the support in a 1:10 volumetric ratio. This cell-laden support was then pipetted into a PDMS-based device and subsequently stabilized via UV crosslinking. On D1, viability appeared to be roughly equivalent in both samples containing channels and those that did not based on the live/dead imaging (Fig. 7 and SI Fig. 5). However, on D4 the cell viability in samples without channels decreased everywhere compared to D1, whereas samples containing channels maintained a region of significantly higher cell viability adjacent to the open channel that extended radially from the channel to 100 µm into the support (Fig. 7A and SI Fig. 5).

**Figure 7:**
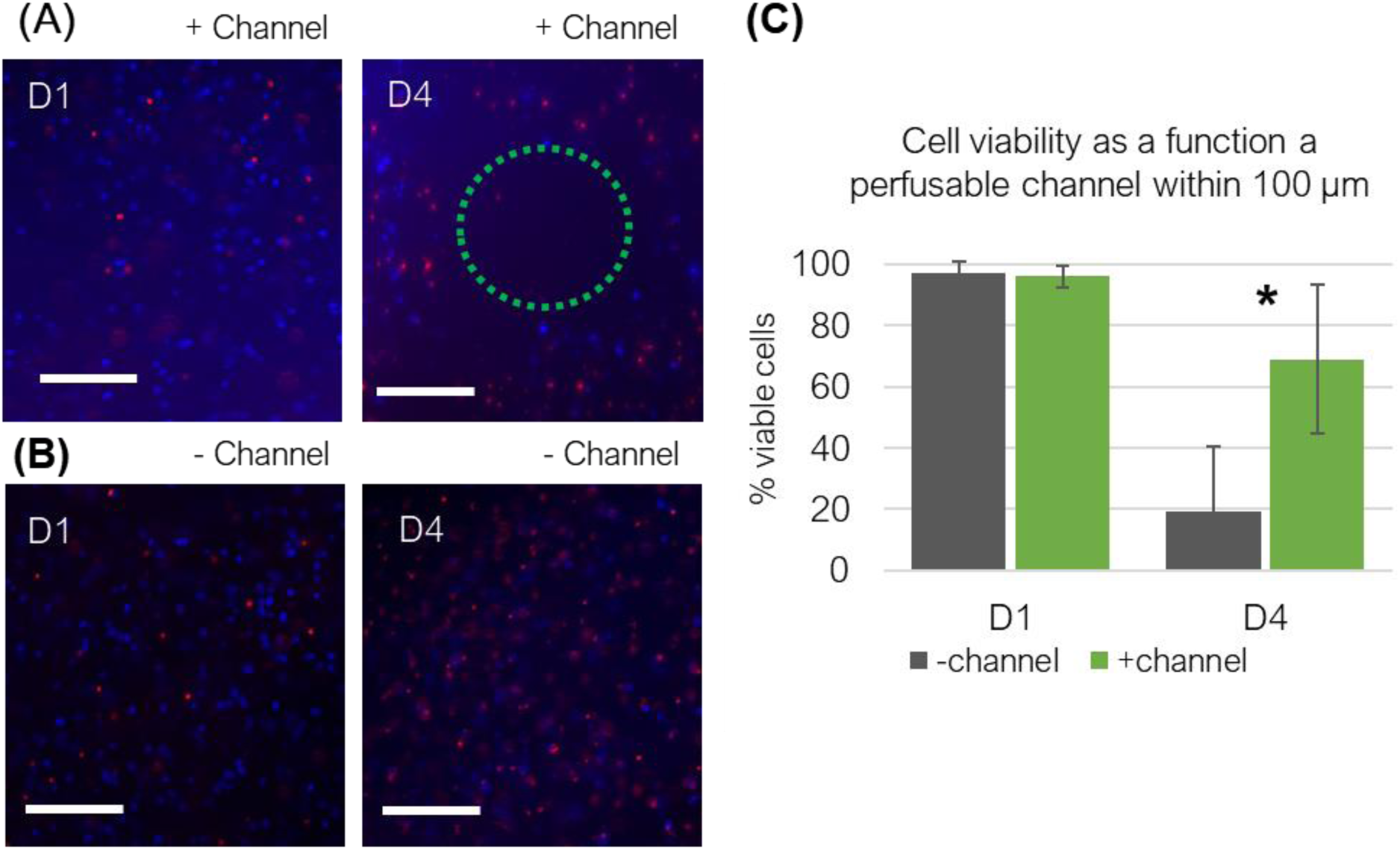
Live/dead imaging of fibroblasts (blue/red dyes, respectively) in cross-sections of (A) perfusable and (B) non-perfusable NorHA supports at days 1 and 4. The green dotted circle in the right panel of (A) indicates the location of the perfusable channel. The area around the perfusable channel shows higher viability compared to any region within the non-perfusable support. (C) Cell viability is similarly in constructs regardless of the presence of a channel on day 1, but is significantly higher adjacent to a printed channel on day 4 in samples that contain channels (* = p < 0.05). Scalebars: 200 µm.

Towards establishing channels that recapitulate the cellular architectures of vessels found within the human body, HUVECs were added to perfusable channels and imaged. HUVECs were found covering all interior surfaces of a 300 µm diameter channel. The top, mid, and bottom planes showed coverage of channel walls by HUVECs on D3 (Fig. 8), after two days of bidirectional flow, which was introduced on D1, 24 h post-HUVEC seeding. Additionally, over time the cells reorganized, and layers lining the lumen showed more continuous and concentrated fluorescence along the surfaces of the channels between D1 and D3 images (Fig. 8D and E, SI Fig. 6).

**Figure 8:**
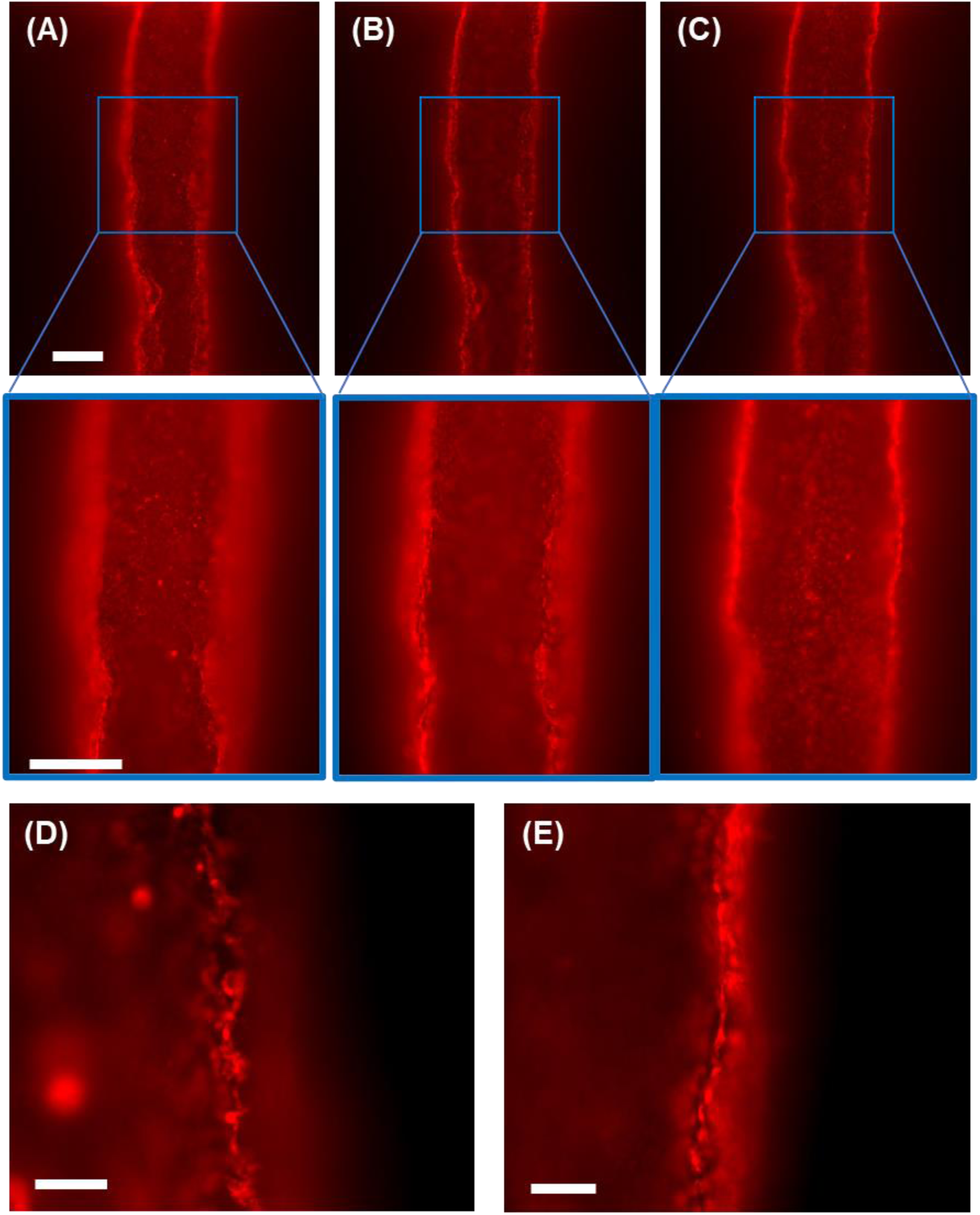
Cels line the surface of an open channel, as seen here using HUVECs labeled with CellTracker Deep Red, with coverage seen in the (A) top plane on D3, (B) mid-plane on D3,and (C)the bottom plane on D3. Cels lining the channel at (D) D1 are shown in close up. (E)A magnified image ofthe lining on D3after2 days of flow reveals changes in morphology. Scalebars: 300 µm.

## 4. Discussion

Biofabrication technologies continue to advance researchers’ capabilities to design cellular and material architectures within engineered tissues and microphysiological systems with increasing complexity and resolution(60). Here, formulating a hydrogel that might be designed to have specific biochemical and biophysical properties(53) as a granular system enables embedded printing to further define material and cellular architectures within the hydrogel. In contrast to processes where reversible embedding is desired – where well-established biomaterial supports include FRESH 2.0(41) or Carbopol(22) – the support material formulated as a granular system here was intended to remain in the final construct. This processing is intended to combine 3D printing processes, which can achieve high-resolution spatial positioning, with small particles to achieve precise definition of cell-containing hydrogel composition(26). When a designable hydrogel biomaterial, such as the NorHA system used here, these powerful biofabrication processes can be combined capabilities to biofunctionalized hydrogels with specific ligands at designed concentrations, with controlled interparticle crosslinking, and with tunable mechanics.

### 4.1 Granular support and ink materials

The NorHA granular support materials formulated here exhibit small average microgel diameters (Fig. 2A-B), approaching that achieved in established gelatin- and poly(acrylic acid)-based granular supports(22,35,41), important to high resolution in embedded printing processes(47). Particle generation is achieved through simple emulsification and results in particle populations with polydisperse size distributions (Fig 2B). Filtration might reduce polydispersity but was not used here.

The rheological responses of all tested granular materials are exhibited behaviors characteristic of other microgel-based materials(44,61). In the unidirectional shear-testing, the tested materials had a yield stress that correlates to particle crosslinking density. The materials were shear-thinning (Fig. 3A), in line with previous work on microgel rheological behavior(22). This shear-thinning behavior was expected as applied stress reached the yield stress of the material, where the particles could freely rearrange resulting in a fluid-like flow of the bulk system.

Rheological frequency sweeps (Fig. 3B) showed that storage moduli for all materials were frequency independent below the yield stress. Storage moduli were dependent on component particle stiffness and were observed to decrease with decreasing crosslinking density as expected. The characteristic minima (Fig. 3B) that were found in the loss modulus curves were a rheological signature of jammed particle systems that result from two events: particle rearrangements occurring at frequencies below the minimum point, and solvent dissipation at frequencies past this point(44). The frequency at which this minimum occurred appeared to correlate to particle crosslinking density: at lower crosslinking densities the minimum was observed at lower frequencies. Since polymer crosslinking density affects viscoelastic properties, this change in frequency could be attributed to an increase in dissipative behavior of the particles, thus leading to a lower frequency when this behavior dominates. Importantly, tests included the interstitial 1% NorHA, and the data indicated that at this concentration the solubilized 1% NorHA did not affect key signature rheological properties, discussed below, of the jammed NorHA microgel supports.

The particle-based materials developed here exhibited typical jammed microgel behavior in response to strain sweeps as well. At low strain, the storage moduli values remained strain-independent and were solid-like, indicated by the higher G’ compared to G”. This was followed by yielding at increasing strain: the G’-G” crossover is observed with a characteristic peak in the loss modulus during yielding. Interestingly, there was a dip in the loss modulus at the onset of yielding but before the modulus crossover for the NorHA-2 material. From a previous study using an emulsion system for printing(62), an overshoot in the low strain regime was observed that appeared to become larger as particle crosslinking density decreased. In that work, this was attributed to contact breaking between particles, but not at a sufficient level to induce flow. This has also been seen elsewhere(63), where a modest overshoot at low strains appeared to be dependent on particle properties. Likely a similar phenomenon was occurring among the softest particles here in the NorHA-2 group. Here, the G’-G” crossover was observed to occur at higher strains as particles became softer. This was attributed to the particles becoming increasingly deformable, resulting in increased packing fraction for soft particles. In other words, softer particles can deform to create more facets with neighboring particles, which in turn increases friction between the particles and would lead to an increase in the G’-G” crossover, as has been reported(63,64).

UV curing of the supports showed a clear, expected increase in G’ of all materials, with increases of roughly 3x-4x. Elastic moduli of crosslinked scaffolds ranged from roughly 500 Pa for NorHA-2 to 1800 Pa for NorHA-8, with moduli of the granular materials increasing with increasing crosslinking density of the component particles, as expected. The range of values obtained here also showed that the support stiffness was tunable and spanned a range of values within reported ranges for soft tissues like brain, lung, small intestine, and liver(65,66).

The rheological properties of the gelatin ink were similar to those of bulk NorHA materials, as expected. The gelatin particles were small and polydisperse, with an average diameter of 4.60 ± 3.77 µm. A smaller diameter was used as particle diameters would ultimately affect resolution of printed features(10,47). In other words, the smaller diameter particles facilitated the printing of finer filaments and features. The ink formulation tested here was shear-thinning, and therefore extrudable through a needle. As in the NorHA materials, a minimum in the loss modulus during frequency sweep testing and a peak in in the strain sweep at the G’-G” crossover that are characteristic of granular systems were observed. Notably, the presence of interstitial HA was – as with the NorHA materials – not observed to affect key rheological properties at concentration used.

### 4.2 3D printed architectures in all-granular hydrogel systems

Filaments of granular hydrogel inks with varying diameters were achieved by varying the printing flow rates. Filaments were printed with diameters as small as 100 µm (Fig. 5A). These small filament diameters relied on the small particle sizes used here and would not have been achievable using particles with sizes typically achieved in microfluidic particle-generation processes, on the order of 100 µm and larger. Continuous and granular support materials, particularly supports comprised of microscale (<100 µm) particles have allowed continuous hydrogel inks to be printed with resolutions below 50 µm in existing work (47). Here, a granular ink comprised of microscale particles could be printed in filaments with diameters on the order of 100 µm and greater, depending on print parameters, which compares favorably with many continuous hydrogel – or hydrogel precursor – inks and bioinks printed into granular materials, where print resolution is often in the mesoscale (100 – 1000 µm) (47). Revisions to print parameters or hardware – for example, smaller diameter nozzles – would be necessary to achieve higher resolution using the granular hydrogel ink formulations employed here.

The use of gelatin microgels allowed filaments to be liquefied under culture conditions, resulting in channels with perfusable lumens whose diameters were slightly smaller than the printed granular filaments. Decreases in final channel diameters are attributed to the use of interstitial NorHA and swelling of the hydrogel support after gelatin ink removal. Perfusable channels had diameters of approximately 200 µm and larger (Fig 5A-B). These channel diameters compare favorably to other 3D printed vessels, where extrusion-based technologies typically can achieve mesoscale vessels with other technology, such as soft lithography, laser-based degellation, and cellular self-assembly typically being required to achieve microscale vascular structures (67).

Perfusion of channels within the fluidic systems we used here limited flow to channels with larger diameters, as hydrostatic pressures were used to drive flow and the magnitudes if these pressures were limited by device dimensions. Approximating the flow the system as a Poiseuille flow, resistance increases exponentially with decreases in channel diameter: a halving of channel diameter results in a 16-fold increase in flow resistance. Due to the retention of fluorescence in the FITC channel (Fig. 5A), there is indication that the lumens of the smallest printed channels may retain the printed gelatin ink, suggesting insufficient pressure to clear these channels. Diffusion of interstitial NorHA from the support into the gelatin ink may also have reduced channel diameter.

The use of perfusable channels to establish concentration gradients is viewed as an important aspect in engineering microphysiological systems – a future application of this technology. Here, we observed diffusion of a small molecule, RhoB, and large protein, FITC-BSA establishing concentration gradients across the granular system with time (Figs. 5D and 5E). The smaller molecule evidenced a higher effective diffusivity, as expected. The gradients within the granular NorHA were seen to be approximately linear, as would also be expected in approximately 1D diffusion. The dip in the concentration RhoB profile within 100 µm, then increase and approximately linear profile from 200 µm onward, evidences a dilution in NorHA concentration – attributed to swelling and loosely crosslinked interstitial NorHA – at the channels’ edges.

Because RhoB and NorHA carry opposing electrostatic charges, RhoB partitions preferentially into regions of higher NorHA concentration, and thus the RhoB concentration profiles observed are consistent with differences in NorHA concentration discussed: a concentration determined by particle design within material bulk, but diluted where intersitial NorHA crosslinked at channel peripheries.

Ultimately, designed placement of channels within a tissue construct with known diffusive characteristics can be used to overcome diffusion limitations typically found in thick tissue constructs, enabling increased mass transport of oxygen and nutrients in cell-laden constructs (67–69). The establishment of gradients using small and large model molecules suggests the ability to include chemoattractants and growth factors to influence cell behaviors or processes such as angiogenesis. Here, transport radially from the channels appears to be largely driven by diffusion. Radial convective perfusion is not evident in experimentation: in visualizing flow with fluorescent microspheres (Fig. 5C), streamlines are in the axial direction, with no microsphere flow radially to the channel or aggregation observed at channel walls, as would be expected if there were radial flows. Perfusion with soluble RITC-dextran (Fig. 5A), with a hydrodynamic radius on the order of 10s of nm (70), also evidences no radial convection from the printed channels. Finally, characterization of diffusion (Fig. 5D), suggests effective diffusivities that are decreased compared to diffusivities in free solution using a rough Stokes-Einstein approximation, where the square distance of solute diffusion is equal to diffusivity times twice the time elapsed. In this simplified analysis, albumin is observed as having an effective diffusivity on the order of 1 × 10^−7^ cm/s^2^, comparable to diffusion rates in nanoporous hydrogels (71), implicating a lack of unhindered interstitial pore spaces for diffusion or convection.

The dominance of diffusive transport in the radial direction is attributed, in this system, to the small hydrogel particle size, preparation of the support material, and the addition of the interstitial NorHA. While overall porosity is dependent on particle packing and polydispersity (72–75), rather than particle size (75), smaller diameter particles will yield smaller pores overall compared to larger diameter microgel particles. Because hydrogels are deformable (72,73), the diffusion-dominated behavior is attributed to the collapse of these small pores under the high centrifugation speeds used in preparing the support material, with the interstitial NorHA helping crosslink the support in this state. Because the granular support hydrogel used here appears to provide a barrier to radial convection, delivery of molecular factors via embedded channels would be expected to depend on a balance between their mass flow rate into the channel and diffusive flow rate radially from the channels, as is the case in other hydrogel systems that support perfusable channels (67–69). The complex structures (Fig. 6) reflect capabilities that will enable printing of architectures or cellular-material features in platforms for tissue engineering. In addition to branching structures found in vasculature, capabilities to depots of printed inks next to filaments (Fig. 6) will facilitate precise establishment of structures such as those seen in studies of stem cell development and tissue engineering where embryoid bodies have been placed next to microfluidic channels(76) or hepatic spheroids being placed between two channels(77). Various types of cancer have also been shown to ablate nearby vascularized channels(78–80) and this could be recapitulated in the system shown here. The system developed here also demonstrated controlled placements of depots of material, enabling arbitrary and complex patterns that are challenging to create with many printing methods. Depots, or voxels (81–83), printed here had diameters of roughly 500 µm, which is comparable to voxels achieved by printing continuous inks into granular supports (81–83) and to depots of continuous or particle inks printed into continuous supports (27,46). Perhaps uniquely to all-particle printing, droplets of an initial granular hydrogel ink could be easily printed within a granular support (Fig. 6C) and a second granular ink could be printed into the initial depot (Fig. 6D). This demonstrated the exciting potential of all-particle based printing to have a printed granular ink function as a support on its own after its deposition, supporting fabrication of complex, multilayered structures.

### 4.3 Introducing cells into support materials and channels in all-granular hydrogel systems

In tissue engineering applications and studies of cellular behaviors in 3D, cells are included within hydrogels. Confirming the potential of the granular matrix to support cell viability, cells included within the NorHA support showed a 10x increase in metabolic activity over nine days compared to D1, when seeded in the lowest density tests of a 1:50 cell suspension volume to hydrogel volume ratio (SI Fig. 4), where the cell suspension contained 100 million fibroblasts/mL. At the highest density tested of 1:10, metabolic activity showed a slight increase of 2x, and at an intermediate density of 1:20, metabolic activity increased approximately 5x. Taken together, these results indicated cell viability is supported within these systems, with a maximum of 10 – 20 million fibroblasts/mL supported under testing conditions of a thin hydrogel in which oxygen and nutrient transport is not limited by hydrogel volume.

By printing channels into thicker NorHA supports containing fibroblasts, we observed both the effects of the printing process and open channels that could facilitate transport on cell viability. Live/dead staining after printing indicated that the process was cytocompatible with respect to embedded cells. In the 4-day culture period following printing, extensive cell death was observed in the constructs that did not contain channels. However, this cell death was mitigated by the inclusion of printed channels. In cross-sections, an area around printed channels showed high cell viability compared to the rest of the construct and the non-perfused construct (Fig. 7 and SI Fig. 5), in line with previous work vascularizing hydrogels(69). This provided evidence that cells were receiving nutrients and oxygen from the printed channels.

Towards achieving endothelialized channels and, ultimately, biomimetic vessel-like structures within all-particle 3D printed systems, we observed that HUVECs were able to line the lumens of printed channels (Fig. 8 and SI Fig. 6) when NorHA microgels were functionalized RGD prior to HUVEC addition. Because flow supports endothelialization of engineered vascular systems(84), flow was introduced to the channels one day after HUVECs were seeded. Images comparing D1, no flow, and D3, after two days of flow, suggested that cells lined the channels after seeding and began to organize into a continuous monolayer along the channel wall after exposure to flow (Fig. 8 and SI Fig. 6). This change in morphology over time would be expected in engineered vascular systems where endothelial cells lining channels mature and adopt morphologies in response to exposure to the dynamic flow environment(59). Further study will be required to determine the effects of material design on endothelial monolayers and to optimize processes for lining the channels. However, analysis of fluorescence due to HUVEC cytoplasmic staining (CellTracker) along channel walls at the midplane of the channel on day 3 post-seeding shows continuous fluorescent signal (SI Fig. 7 A and B). Additionally, microscopy images from focal planes near the top walls of channels show cobblestone-like cellular morphologies covering the channels’ top surfaces (SI Fig. 7 C).

As cells can be included within the granular support hydrogel or in the printed material to create bioinks, the system offers capabilities to fabricate complex multicellular structures. Multilayered structures have applications in modeling certain tissue types or tumors. Cell signaling gradients might ultimately be created by careful positioning of multiple cell populations relative to one another. Bioinks containing either HUVECs or mesenchymal stromal cells were printed one into the other in a depot-into-depot structure (SI Fig. 8A) or a filament-intesecting-filament arrangement (SI Fig. 8B). Additionally, channels that can be lined with HUVECs can be established directly within cell-containing granular support materials (SI Fig. 8C and D). These biofabrication capabilities offer avenues to increasing complexity and biomimicry in bioprinted tissue and microphysiological systems.

## 5. Conclusions

The combination of bioprinting and granular hydrogel technologies offer new possibilities for designing and fabricating complex hydrogel and engineered tissue structures. In particular, the work here demonstrates how these technologies might be used towards achieving the heterogeneity of biological structures that feature complex physical and tethered biochemical cues, soluble gradients of cell-instructive compounds, and vascular structures. Key to this approach were a highly tunable granular NorHA support matrix and a removable granular ink. Both the support and ink biomaterials were based on small hydrogel microparticles formed by emulsification and designed to have material properties compatible with the bioprinting process. High resolution features and unique structures, such as inks deposited within inks, were possible through the combined use of granular supports and inks. Cells included within the support material are compatible with the bioprinting processes, and channels crucial to alleviating transport limitations could be printed. Endothelial cells introduced into these channels could adhere to their luminal surfaces and developed as linings with flow. This study demonstrates the basis of this all-granular bioprinting system for establishing a 3D canvas that might be extended towards creating complex, cell-laden materials for biomedical applications.

## Acknowledgements

This work was funded by the National Science Foundation through Future Manufacturing Seed grant 2036968 and the National Institute of General Medical Sciences at the National Institutes of Health through grant R35GM147410.

## Supplemental information

**SI Table 1:** Average particle size and CV.

| Material | Diameter ($\mu\text{m}$) | CV (%) | n |
| --- | --- | --- | --- |
| NorHA-8 | $9.63 \pm 4.50$ | 46.7 | 393 |
| NorHA-4 | $16.44 \pm 10.47$ | 63.7 | 466 |
| NorHA-2 | $16.45 \pm 9.81$ | 59.6 | 427 |
| Gelatin Ink | $4.60 \pm 3.77$ | 82.0 | 520 |

**SI Table 2:**
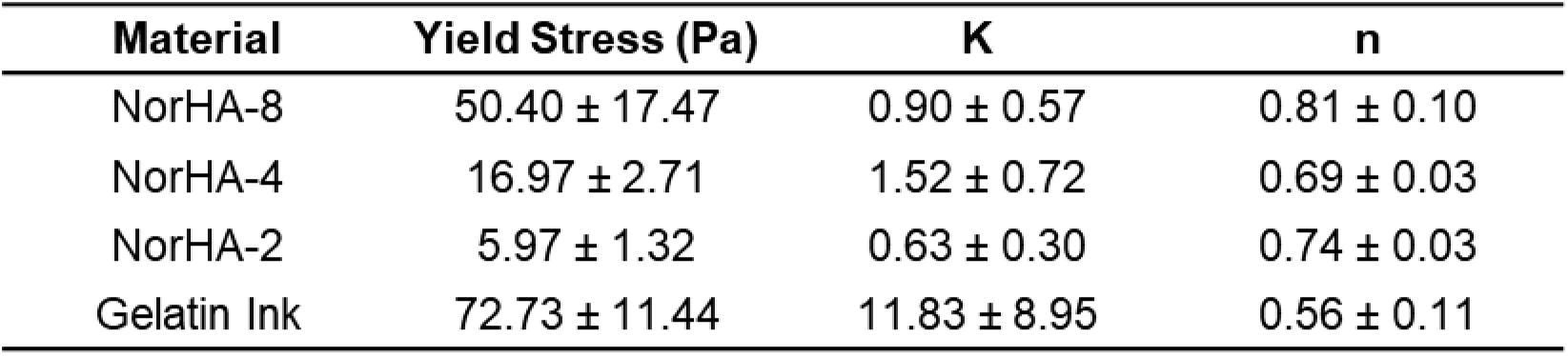
Herschel-Bulkley parameters.

**SI Table 3:**
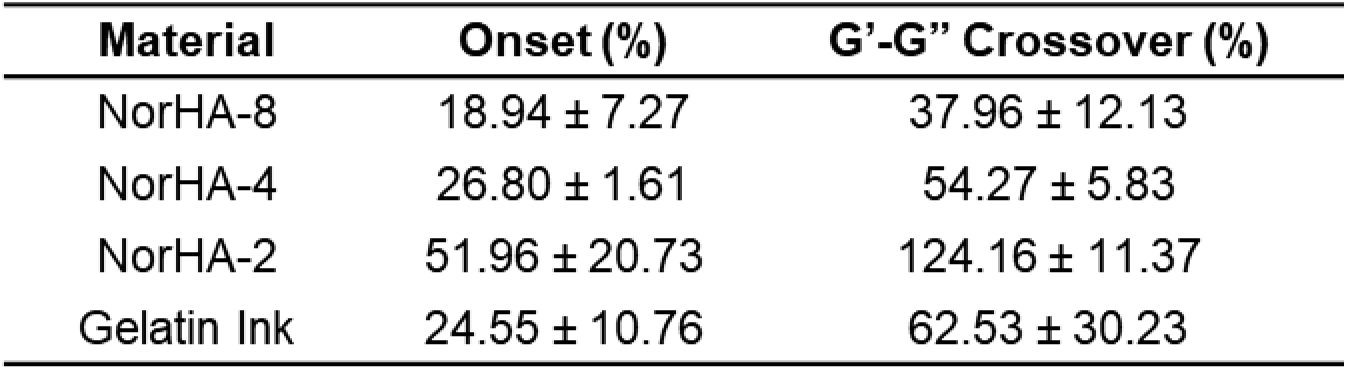
Strain sweep onset and G’-G" crossover.

| Material | Onset (%) | $G'$-$G''$ Crossover (%) |
| --- | --- | --- |
| NorHA-8 | $18.94 \pm 7.27$ | $37.96 \pm 12.13$ |
| NorHA-4 | $26.80 \pm 1.61$ | $54.27 \pm 5.83$ |
| NorHA-2 | $51.96 \pm 20.73$ | $124.16 \pm 11.37$ |
| Gelatin Ink | $24.55 \pm 10.76$ | $62.53 \pm 30.23$ |

**SI Fig. 1:**
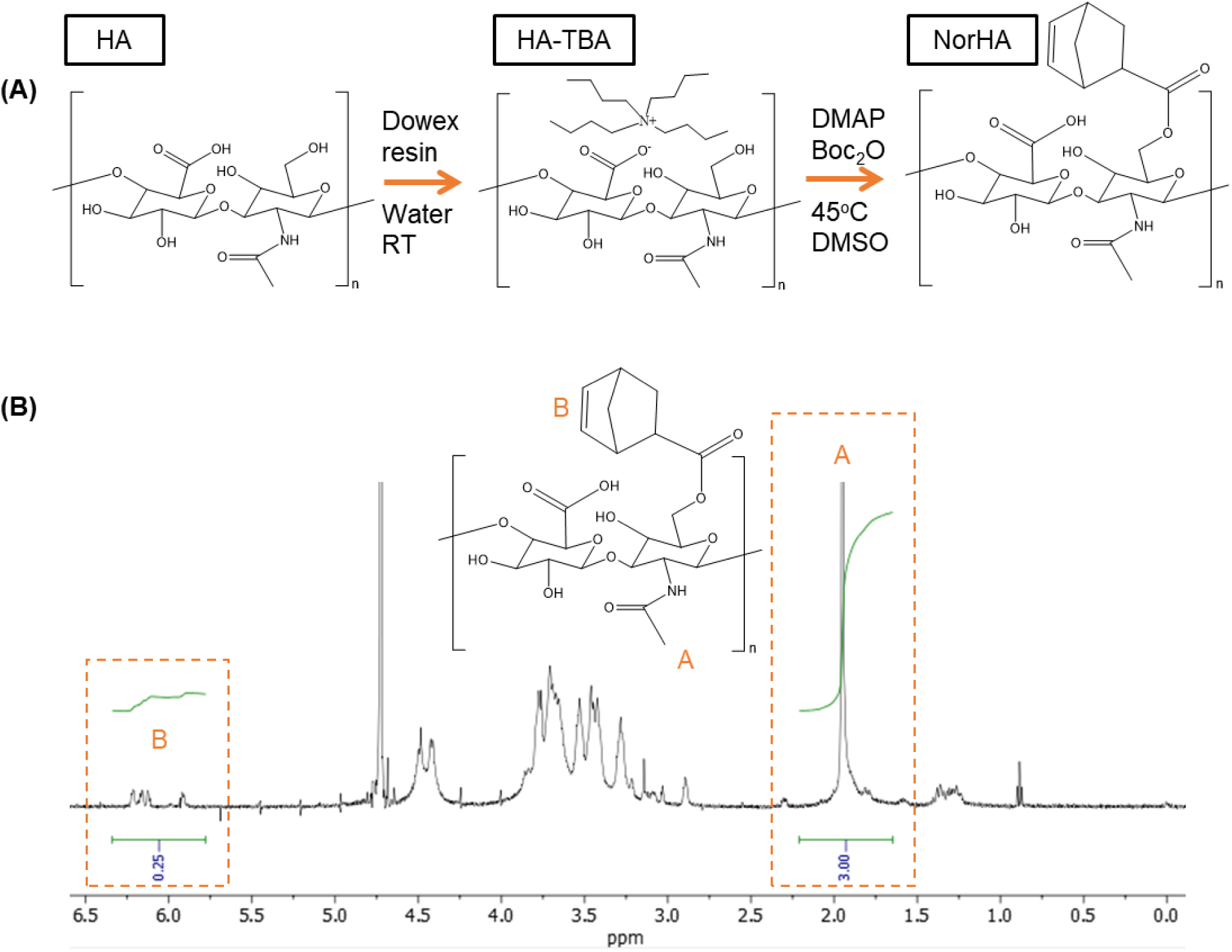
Synthesis and NMR. (A) synthesis of NorHA using a multistep process, and (B) an example of NMR spectrum for NorHA, showing roughly 12.5% modification.

**SI Fig. 2:**
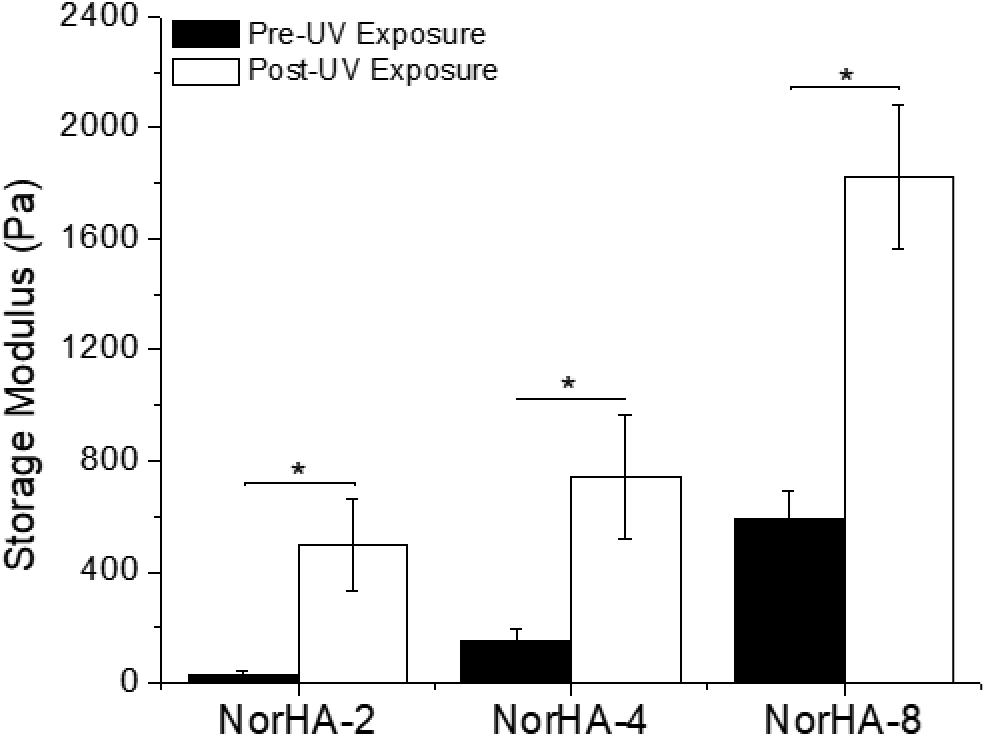
G’ before and after UV crosslinking. Average storage modulus before and aftercrosslinking of NorHA-2, -4, and - 8 microgels with interstitial NorHA. As expected, the storage modulus is significantly different post-crosslinking, and is dependent on crosslinking density of the particles themselves.

**SI Fig. 3:**
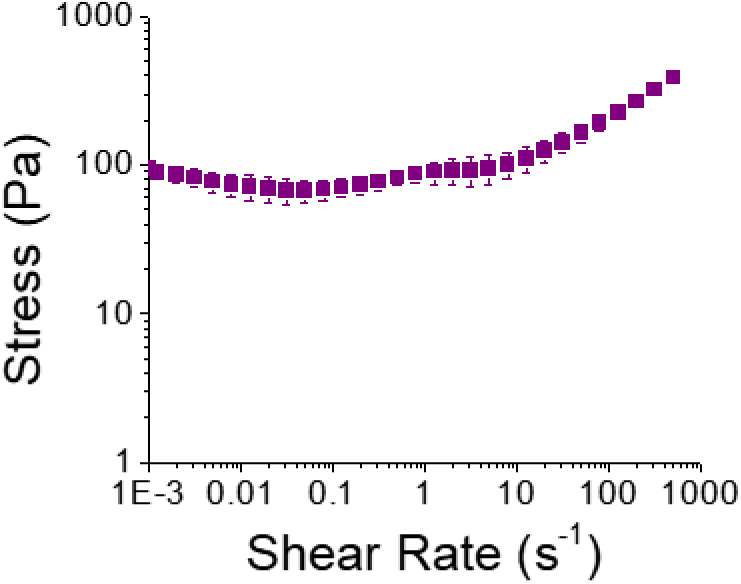
Gelatin Ink yield stress. Yield stresstesting of gelatin microgel-based ink, with error bars denoting standard deviation.

**SI Fig. 4:**
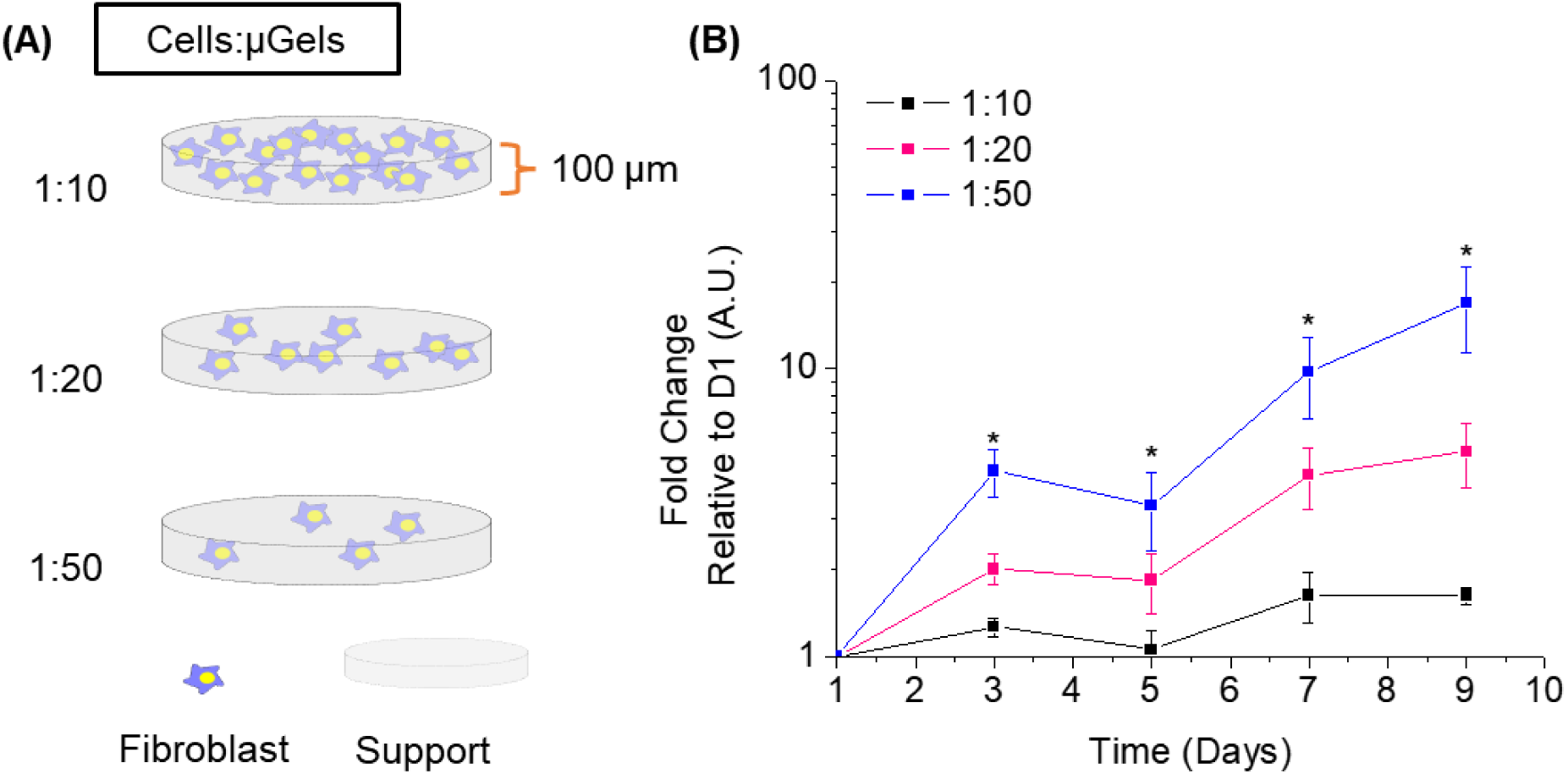
Cell proliferation in thin gels as a function of cells: support. (A)fibroblast proliferation experiment setup with varying eel densities within NorHA-4 supports, and (B) Alamar blue assay results over a 9-day time period showing either maintenance of eel viability or proliferation of fibroblasts within the NorHA-4supports_

**SI Fig. 5:**
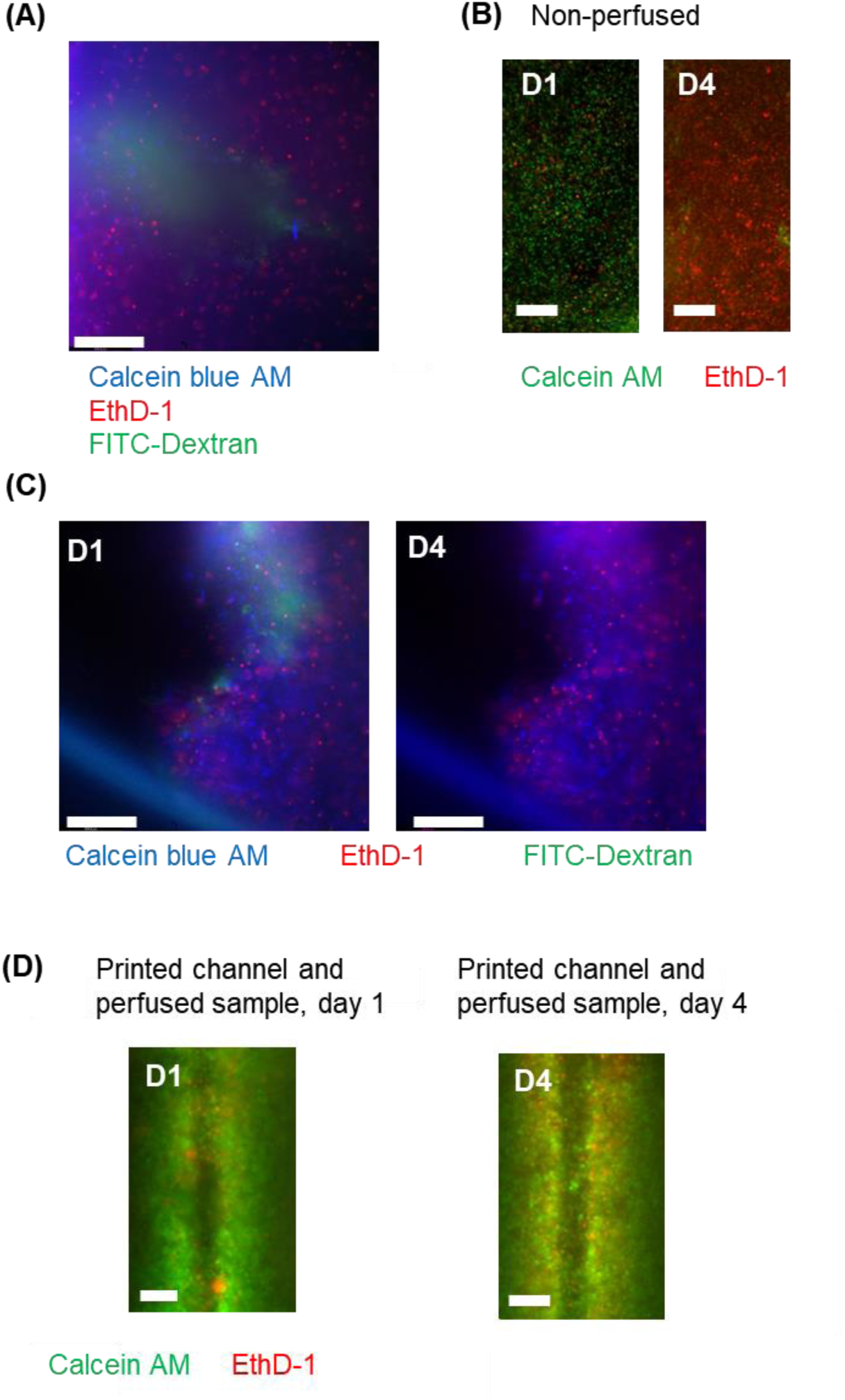
Cell viability in the presences of channels. (A) Channel from Fig. 7 but with a FITC introduced into the channel lumen, (B) another non­ perfused sample ilustrating low viability from D1to D4as expected, (C)Additional samples with cross­ sectioned channels on D1 and D4, (D)longitudinal plane of channel on D1 and D4. Scalebars for al images are 300µm.

**SI Fig. 6:**
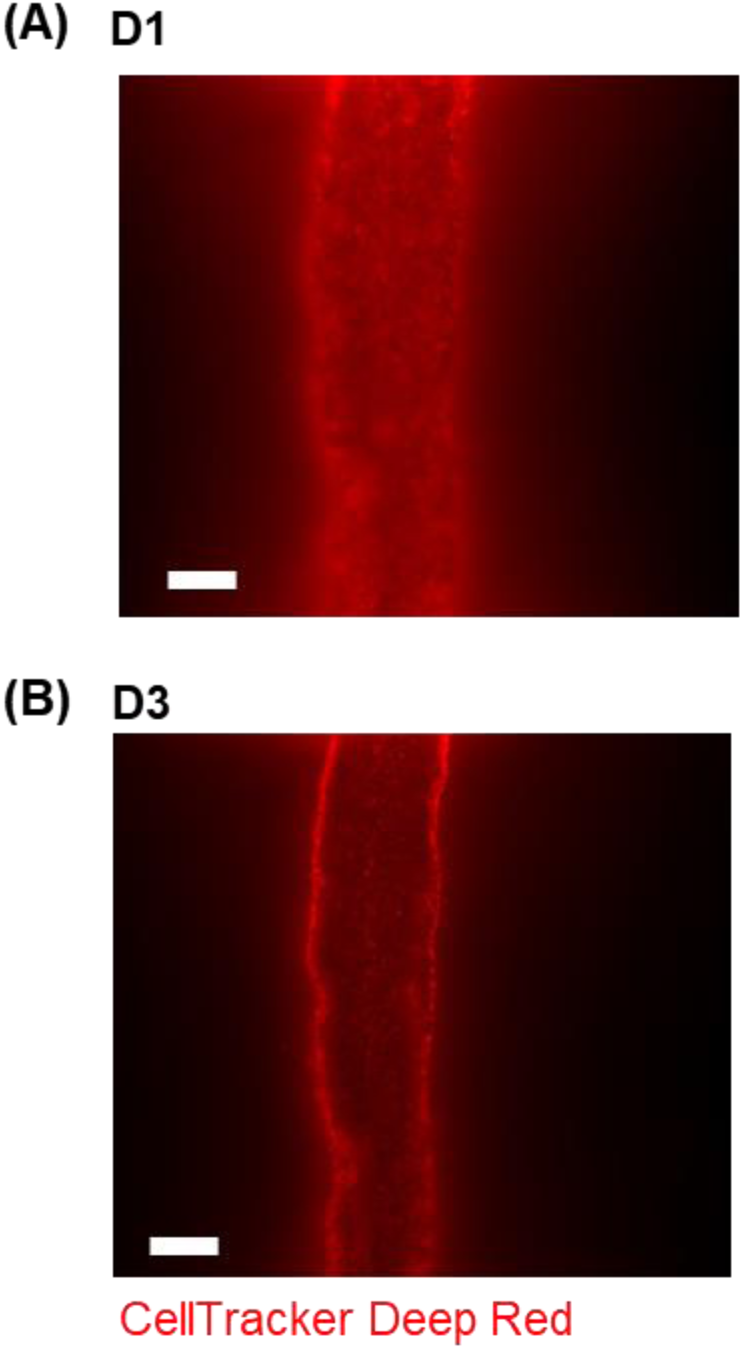
Endothelial cells on channel walls - day 1 vs day 3. Images of channel lined with HUVECs on (A) D1 post-seeding, and (B) D3 post­ seeding. Scalebars: 300 µm.

**SI Fig. 7:**
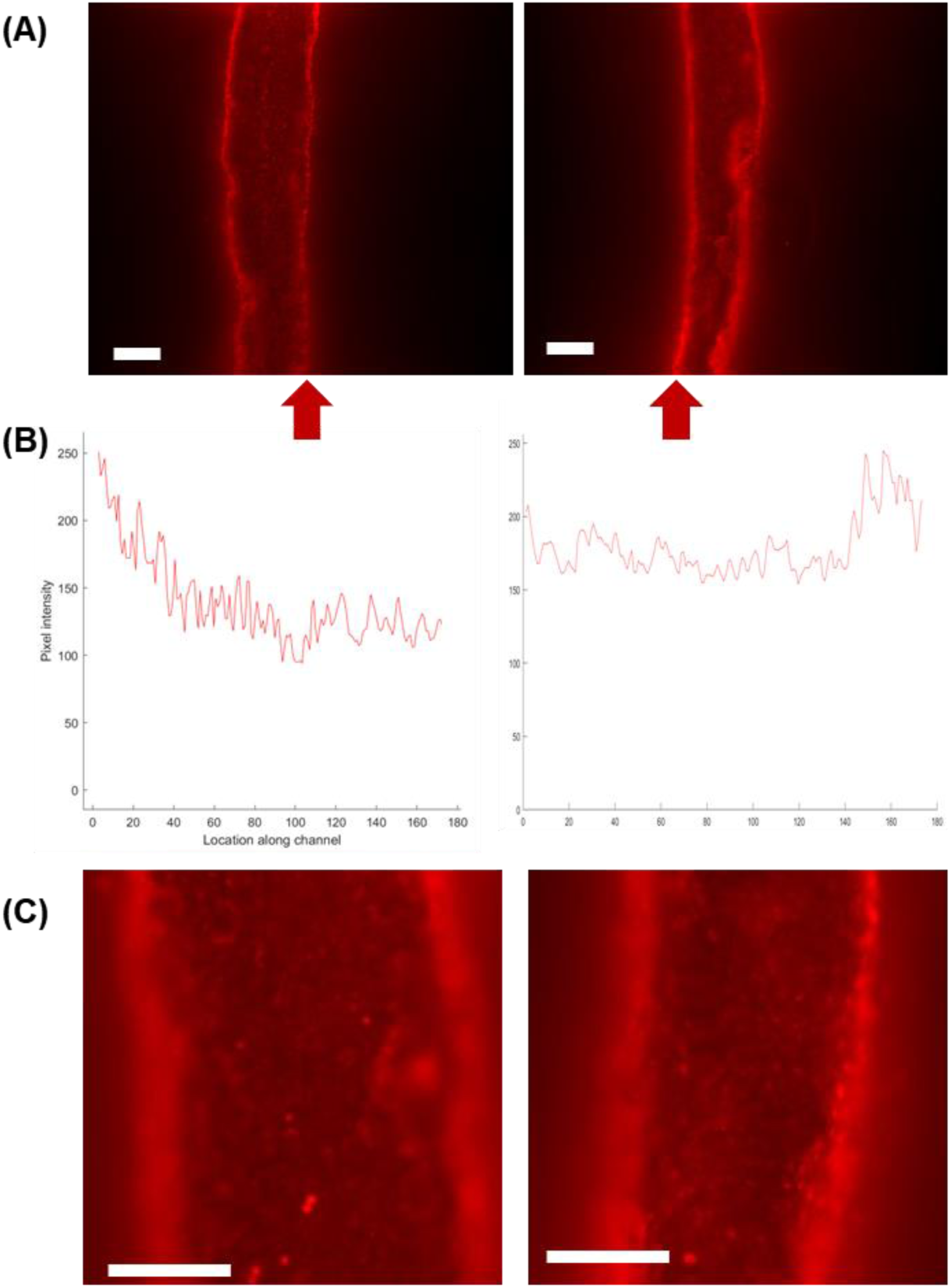
Analysis of endothelial cell coverage on chennel walls on days 3. (A) Images of channels lined with HUVECs on D3 post-seeding. straight walls, indicated by arrows. used in analysis of coverage. (B) Quantification. using MATLAB’s improfile function, of fluorescent intensity along channel wals due to HUVECs’ CellTracker dye. (C) Images from focal planes near top surfacesof channels showing HUVEC coverage. Scalebars: 300µm (A) and 150 µm (B).

**SI Fig 8:**
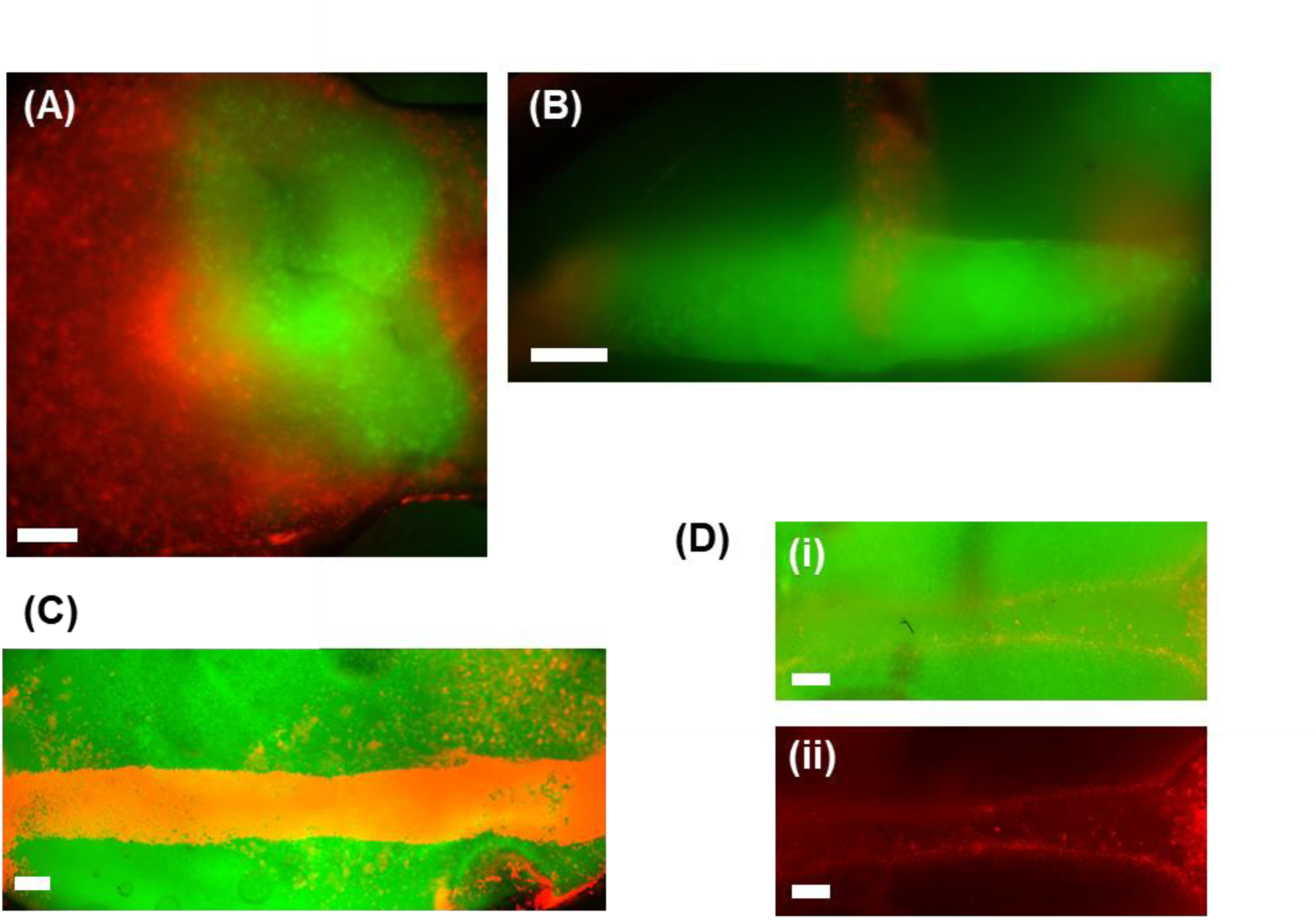
Printing multiple bioinks into complex Printing multiple structures. granular bioinks into complex structures in a granular support material_ In **al** images, HUVECs are labeled red and mesenchymal stromal eels (MSCs) are green)_ A) MSC depot is printed into a HUVEC depot to create a layered structure_ B) HUVEC-containing filament (- 400µm long) printed orthogonaly at midpoint of a longer MSC-containing filament C) A channel is printed into an MSC-containing support material into which HUVECs are introduced towards creating an endothelialized channel within a hydrogel containing another eel population_ Image taken immediately after introducing HUVECs_ D) The same channel from C, 2 hours later after washing out unadhered HUVECs_ i) Image with the HUVEC (red)and MSC (green) fluorescent channels overlaid_ ii) Image of HUVECs only show eel-lined channel wals at 2 hours post-HUVEC seeding_ Scalebars: 200µm_

## References

1. Moroni L, Burdick JA, Highley C, Lee SJ, Morimoto Y, Takeuchi S, et al. Biofabrication strategies for 3D in vitro models and regenerative medicine. Nat Rev Mater. 2018 May;3(5):21–37.

2. Chung JJ, Im H, Kim SH, Park JW, Jung Y. Toward Biomimetic Scaffolds for Tissue Engineering: 3D Printing Techniques in Regenerative Medicine. Front Bioeng Biotechnol [Internet]. 2020 Nov 4 [cited 2024 May 3];8. Available from: https://www.frontiersin.org/articles/10.3389/fbioe.2020.586406

3. Lee VK, Dai G. Printing of Three-Dimensional Tissue Analogs for Regenerative Medicine. Ann Biomed Eng. 2017 Jan 1;45(1):115–31.

4. Celikkin N, Mastrogiacomo S, Dou W, Heerschap A, Oosterwijk E, Walboomers XF, et al. In vitro and in vivo assessment of a 3D printable gelatin methacrylate hydrogel for bone regeneration applications. Journal of Biomedical Materials Research Part B: Applied Biomaterials. 2022;110(9):2133–45.

5. Yan Y, Chen H, Zhang H, Guo C, Yang K, Chen K, et al. Vascularized 3D printed scaffolds for promoting bone regeneration. Biomaterials. 2019 Jan 1;190–191:97–110.

6. Zhang M, Qian T, Deng Z, Hang F. 3D printed double-network alginate hydrogels containing polyphosphate for bioenergetics and bone regeneration. International Journal of Biological Macromolecules. 2021 Oct 1;188:639–48.

7. Zhang X, Zhang H, Zhang Y, Huangfu H, Yang Y, Qin Q, et al. 3D printed reduced graphene oxide-GelMA hybrid hydrogel scaffolds for potential neuralized bone regeneration. J Mater Chem B. 2023 Feb 8;11(6):1288–301.

8. Kang HW, Lee SJ, Ko IK, Kengla C, Yoo JJ, Atala A. A 3D bioprinting system to produce human-scale tissue constructs with structural integrity. Nat Biotechnol. 2016 Mar;34(3):312– 9.

9. Choi YJ, Kim TG, Jeong J, Yi HG, Park JW, Hwang W, et al. 3D Cell Printing of Functional Skeletal Muscle Constructs Using Skeletal Muscle-Derived Bioink. Advanced Healthcare Materials. 2016;5(20):2636–45.

10. Skylar-Scott MA, Uzel SGM, Nam LL, Ahrens JH, Truby RL, Damaraju S, et al. Biomanufacturing of organ-specific tissues with high cellular density and embedded vascular channels. Science Advances. 2019 Sep 6;5(9):eaaw2459.

11. Han H, Park Y, Choi Y mi, Yong U, Kang B, Shin W, et al. A Bioprinted Tubular Intestine Model Using a Colon-Specific Extracellular Matrix Bioink. Advanced Healthcare Materials. 2022;11(2):2101768.

12. Kim W, Kim GH. An intestinal model with a finger-like villus structure fabricated using a bioprinting process and collagen/SIS-based cell-laden bioink. Theranostics. 2020 Jan 22;10(6):2495–508.

13. Kim W, Kim G. Intestinal Villi Model with Blood Capillaries Fabricated Using Collagen-Based Bioink and Dual-Cell-Printing Process. ACS Appl Mater Interfaces. 2018 Dec 5;10(48):41185–96.

14. Heichel DL, Tumbic JA, Boch ME, Ma AWK, Burke KA. Silk fibroin reactive inks for 3D printing crypt-like structures. Biomed Mater. 2020 Aug;15(5):055037.

15. Torras N, Zabalo J, Abril E, Carré A, García-Díaz M, Martínez E. A bioprinted 3D gut model with crypt-villus structures to mimic the intestinal epithelial-stromal microenvironment. Biomaterials Advances. 2023 Oct 1;153:213534.

16. Taebnia N, Zhang R, Kromann EB, Dolatshahi-Pirouz A, Andresen TL, Larsen NB. Dual-Material 3D-Printed Intestinal Model Devices with Integrated Villi-like Scaffolds. ACS Appl Mater Interfaces. 2021 Dec 15;13(49):58434–46.

17. Song Y, Su X, Firouzian KF, Fang Y, Zhang T, Sun W. Engineering of brain-like tissue constructs via 3D Cell-printing technology. Biofabrication. 2020 May;12(3):035016.

18. Brassard JA, Nikolaev M, Hübscher T, Hofer M, Lutolf MP. Recapitulating macro-scale tissue self-organization through organoid bioprinting. Nat Mater. 2021 Jan;20(1):22–9.

19. Negro A, Cherbuin T, Lutolf MP. 3D Inkjet Printing of Complex, Cell-Laden Hydrogel Structures. Sci Rep. 2018 Nov 20;8(1):17099.

20. McCormack A, Highley CB, Leslie NR, Melchels FPW. 3D Printing in Suspension Baths: Keeping the Promises of Bioprinting Afloat. Trends in Biotechnology. 2020 Jun 1;38(6):584– 93.

21. Brunel LG, Hull SM, Heilshorn SC. Engineered assistive materials for 3D bioprinting: support baths and sacrificial inks. Biofabrication. 2022 May;14(3):032001.

22. Bhattacharjee T, Zehnder SM, Rowe KG, Jain S, Nixon RM, Sawyer WG, et al. Writing in the granular gel medium. Science Advances. 2015 Sep 25;1(8):e1500655.

23. Hinton TJ, Jallerat Q, Palchesko RN, Park JH, Grodzicki MS, Shue HJ, et al. Three-dimensional printing of complex biological structures by freeform reversible embedding of suspended hydrogels. Science Advances. 2015 Oct 23;1(9):e1500758.

24. Wu W, DeConinck A, Lewis JA. Omnidirectional Printing of 3D Microvascular Networks. Advanced Materials. 2011;23(24):H178–83.

25. Jin Y, Compaan A, Bhattacharjee T, Huang Y. Granular gel support-enabled extrusion of three-dimensional alginate and cellular structures. Biofabrication. 2016 Jun;8(2):025016.

26. Bhattacharjee T, Gil CJ, Marshall SL, Urueña JM, O’Bryan CS, Carstens M, et al. Liquid-like Solids Support Cells in 3D. ACS Biomater Sci Eng. 2016 Oct 10;2(10):1787–95.

27. Highley CB, Rodell CB, Burdick JA. Direct 3D Printing of Shear-Thinning Hydrogels into Self-Healing Hydrogels. Advanced Materials. 2015;27(34):5075–9.

28. Zhu J, He Y, Wang Y, Cai LH. Voxelated bioprinting of modular double-network bio-ink droplets [Internet]. bioRxiv; 2023 [cited 2024 May 2]. p. 2023.09.19.558463. Available from: https://www.biorxiv.org/content/10.1101/2023.09.19.558463v1

29. Compaan AM, Song K, Chai W, Huang Y. Cross-Linkable Microgel Composite Matrix Bath for Embedded Bioprinting of Perfusable Tissue Constructs and Sculpting of Solid Objects. ACS Appl Mater Interfaces. 2020 Feb 19;12(7):7855–68.

30. Song KH, Highley CB, Rouff A, Burdick JA. Complex 3D-Printed Microchannels within Cell-Degradable Hydrogels. Advanced Functional Materials. 2018;28(31):1801331.

31. Kajtez J, Wesseler MF, Birtele M, Khorasgani FR, Rylander Ottosson D, Heiskanen A, et al. Embedded 3D Printing in Self-Healing Annealable Composites for Precise Patterning of Functionally Mature Human Neural Constructs. Advanced Science. 2022;9(25):2201392.

32. Bakht SM, Gomez-Florit M, Lamers T, Reis RL, Domingues RMA, Gomes ME. 3D Bioprinting of Miniaturized Tissues Embedded in Self-Assembled Nanoparticle-Based Fibrillar Platforms. Advanced Functional Materials. 2021;31(46):2104245.

33. Trikalitis VD, Kroese NJJ, Kaya M, Cofiño-Fabres C, Den S ten, Khalil ISM, et al. Embedded 3D printing of dilute particle suspensions into dense complex tissue fibers using shear thinning xanthan baths. Biofabrication. 2022 Dec;15(1):015014.

34. Abaci A, Guvendiren M. 3D bioprinting of dense cellular structures within hydrogels with spatially controlled heterogeneity. Biofabrication. 2024 Jun;16(3):035027.

35. Morley CD, Ellison ST, Bhattacharjee T, O’Bryan CS, Zhang Y, Smith KF, et al. Quantitative characterization of 3D bioprinted structural elements under cell generated forces. Nat Commun. 2019 Jul 10;10(1):3029.

36. Patrício SG, Sousa LR, Correia TR, Gaspar VM, Pires LS, Luís JL, et al. Freeform 3D printing using a continuous viscoelastic supporting matrix. Biofabrication. 2020 May;12(3):035017.

37. Monteiro RF, Bakht SM, Gomez-Florit M, Stievani FC, Alves ALG, Reis RL, et al. Writing 3D In Vitro Models of Human Tendon within a Biomimetic Fibrillar Support Platform. ACS Appl Mater Interfaces. 2023 Nov 8;15(44):50598–611.

38. Seymour AJ, Westerfield AD, Cornelius VC, Skylar-Scott MA, Heilshorn SC. Bioprinted microvasculature: progressing from structure to function. Biofabrication. 2022 Feb;14(2):022002.

39. Malda J, Visser J, Melchels FP, Jüngst T, Hennink WE, Dhert WJA, et al. 25th Anniversary Article: Engineering Hydrogels for Biofabrication. Advanced Materials. 2013;25(36):5011– 28.

40. Özgen Öztürk-Öncel M, Hiram Leal-Martínez B. F., Monteiro R E., Gomes M A., Domingues RM. A dive into the bath: embedded 3D bioprinting of freeform in vitro models. Biomaterials Science. 2023;11(16):5462–73.

41. Lee A, Hudson AR, Shiwarski DJ, Tashman JW, Hinton TJ, Yerneni S, et al. 3D bioprinting of collagen to rebuild components of the human heart. Science. 2019 Aug 2;365(6452):482– 7.

42. de Gennes PG. Granular matter: a tentative view. Rev Mod Phys. 1999 Mar 1;71(2):S374– 82.

43. Liu AJ, Nagel SR. Jamming is not just cool any more. Nature. 1998 Nov;396(6706):21–2.

44. Menut P, Seiffert S, Sprakel J, Weitz DA. Does size matter? Elasticity of compressed suspensions of colloidal- and granular-scale microgels. Soft Matter. 2011 Dec 1;8(1):156– 64.

45. Divoux T, Tamarii D, Barentin C, Manneville S. Transient Shear Banding in a Simple Yield Stress Fluid. Phys Rev Lett. 2010 May 18;104(20):208301.

46. Highley CB, Song KH, Daly AC, Burdick JA. Jammed Microgel Inks for 3D Printing Applications. Advanced Science. 2019;6(1):1801076.

47. Xie ZT, Kang DH, Matsusaki M. Resolution of 3D bioprinting inside bulk gel and granular gel baths. Soft Matter. 2021;17(39):8769–85.

48. Daly AC. Granular Hydrogels in Biofabrication: Recent Advances and Future Perspectives. Advanced Healthcare Materials. n/a(n/a):2301388.

49. Griffin DR, Weaver WM, Scumpia P, Di Carlo D, Segura T. Accelerated wound healing by injectable microporous gel scaffolds assembled from annealed building blocks. Nat Mater. 2015 Jul;14(7):737–44.

50. Muir VG, Qazi TH, Weintraub S, Torres Maldonado BO, Arratia PE, Burdick JA. Sticking Together: Injectable Granular Hydrogels with Increased Functionality via Dynamic Covalent Inter-Particle Crosslinking. Small. 2022;18(36):2201115.

51. Muir VG, Weintraub S, Dhand AP, Fallahi H, Han L, Burdick JA. Influence of Microgel and Interstitial Matrix Compositions on Granular Hydrogel Composite Properties. Advanced Science. 2023;10(10):2206117.

52. Roh S, Parekh DP, Bharti B, Stoyanov SD, Velev OD. 3D Printing by Multiphase Silicone/Water Capillary Inks. Advanced Materials. 2017;29(30):1701554.

53. Gramlich WM, Kim IL, Burdick JA. Synthesis and orthogonal photopatterning of hyaluronic acid hydrogels with thiol-norbornene chemistry. Biomaterials. 2013 Dec 1;34(38):9803–11.

54. Wade RJ, Bassin EJ, Gramlich WM, Burdick JA. Nanofibrous Hydrogels with Spatially Patterned Biochemical Signals to Control Cell Behavior. Advanced Materials. 2015;27(8):1356–62.

55. Lee W, Lee V, Polio S, Keegan P, Lee JH, Fischer K, et al. On-demand three-dimensional freeform fabrication of multi-layered hydrogel scaffold with fluidic channels. Biotechnology and Bioengineering. 2010;105(6):1178–86.

56. Seymour AJ, Shin S, Heilshorn SC. 3D Printing of Microgel Scaffolds with Tunable Void Fraction to Promote Cell Infiltration. Advanced Healthcare Materials. 2021;10(18):2100644.

57. Wu J, Wang R, Yu H, Li G, Xu K, C. Tien N, et al. Inkjet-printed microelectrodes on PDMS as biosensors for functionalized microfluidic systems. Lab on a Chip. 2015;15(3):690–5.

58. Reid JA, Mollica PA, Johnson GD, Ogle RC, Bruno RD, Sachs PC. Accessible bioprinting: adaptation of a low-cost 3D-printer for precise cell placement and stem cell differentiation. Biofabrication. 2016 Jun;8(2):025017.

59. Kinstlinger IS, Calderon GA, Royse MK, Means AK, Grigoryan B, Miller JS. Perfusion and endothelialization of engineered tissues with patterned vascular networks. Nat Protoc. 2021 Jun;16(6):3089–113.

60. Zandrini T, Florczak S, Levato R, Ovsianikov A. Breaking the resolution limits of 3D bioprinting: future opportunities and present challenges. Trends in Biotechnology. 2023 May 1;41(5):604–14.

61. O’Bryan CS, Kabb CP, Sumerlin BS, Angelini TE. Jammed Polyelectrolyte Microgels for 3D Cell Culture Applications: Rheological Behavior with Added Salts. ACS Applied Bio Materials [Internet]. 2019 Feb 26 [cited 2024 Mar 7]; Available from: https://pubs.acs.org/doi/full/10.1021/acsabm.8b00784

62. Sridharan S, Meinders MBJ, Sagis LM, Bitter JH, Nikiforidis CV. Jammed Emulsions with Adhesive Pea Protein Particles for Elastoplastic Edible 3D Printed Materials. Advanced Functional Materials. 2021;31(45):2101749.

63. Hirsch M, Charlet A, Amstad E. 3D Printing of Strong and Tough Double Network Granular Hydrogels. Advanced Functional Materials. 2021;31(5):2005929.

64. Emiroglu DB, Bekcic A, Dranseike D, Zhang X, Zambelli T, deMello AJ, et al. Building block properties govern granular hydrogel mechanics through contact deformations. Science Advances. 2022 Dec 16;8(50):eadd8570.

65. Chaudhuri O, Gu L, Klumpers D, Darnell M, Bencherif SA, Weaver JC, et al. Hydrogels with tunable stress relaxation regulate stem cell fate and activity. Nature Mater. 2016 Mar;15(3):326–34.

66. Lou L, Paolino L, Agarwal A. Bridging the Gap in Ashby’s Map for Soft Material Properties for Tissue Engineering. ACS Appl Mater Interfaces. 2023 May 24;15(20):24197–208.

67. Fleischer S, Tavakol DN, Vunjak-Novakovic G. From Arteries to Capillaries: Approaches to Engineering Human Vasculature. Advanced Functional Materials. 2020;30(37):1910811.

68. Choi NW, Cabodi M, Held B, Gleghorn JP, Bonassar LJ, Stroock AD. Microfluidic scaffolds for tissue engineering. Nature Materials. 2007 Nov;6(11):908–15.

69. Miller JS, Stevens KR, Yang MT, Baker BM, Nguyen DHT, Cohen DM, et al. Rapid casting of patterned vascular networks for perfusable engineered 3D tissues. Nat Mater. 2012 Sep;11(9):768–74.

70. Armstrong JK, Wenby RB, Meiselman HJ, Fisher TC. The Hydrodynamic Radii of Macromolecules and Their Effect on Red Blood Cell Aggregation. Biophys J. 2004 Dec;87(6):4259–70.

71. Liang S, Xu J, Weng L, Dai H, Zhang X, Zhang L. Protein diffusion in agarose hydrogel in situ measured by improved refractive index method. Journal of Controlled Release. 2006 Oct 10;115(2):189–96.

72. Riley L, Schirmer L, Segura T. Granular hydrogels: emergent properties of jammed hydrogel microparticles and their applications in tissue repair and regeneration. Current Opinion in Biotechnology. 2019 Dec 1;60:1–8.

73. Daly AC, Riley L, Segura T, Burdick JA. Hydrogel microparticles for biomedical applications. Nature Reviews Materials. 2020 Jan;5(1):20–43.

74. Qazi TH, Muir VG, Burdick JA. Methods to Characterize Granular Hydrogel Rheological Properties, Porosity, and Cell Invasion. ACS Biomater Sci Eng. 2022 Apr 11;8(4):1427–42.

75. Riley L, Wei G, Bao Y, Cheng P, Wilson KL, Liu Y, et al. Void Volume Fraction of Granular Scaffolds. Small. 2023;19(40):2303466.

76. Clark AT. Human embryo implantation modelled in microfluidic channels. Nature. 2019 Sep;573(7774):350–1.

77. Bonanini F, Kurek D, Previdi S, Nicolas A, Hendriks D, de Ruiter S, et al. In vitro grafting of hepatic spheroids and organoids on a microfluidic vascular bed. Angiogenesis. 2022 Nov 1;25(4):455–70.

78. Watkins S, Robel S, Kimbrough IF, Robert SM, Ellis-Davies G, Sontheimer H. Disruption of astrocyte–vascular coupling and the blood–brain barrier by invading glioma cells. Nat Commun. 2014 Jun 19;5(1):4196.

79. Silvestri VL, Henriet E, Linville RM, Wong AD, Searson PC, Ewald AJ. A Tissue-Engineered 3D Microvessel Model Reveals the Dynamics of Mosaic Vessel Formation in Breast Cancer. Cancer Research. 2020 Oct 2;80(19):4288–301.

80. Nguyen DHT, Lee E, Alimperti S, Norgard RJ, Wong A, Lee JJK, et al. A biomimetic pancreatic cancer on-chip reveals endothelial ablation via ALK7 signaling. Science Advances. 2019 Aug 28;5(8):eaav6789.

81. Zhu J, He Y, Kong L, He Z, Kang KY, Grady SP, et al. Digital Assembly of Spherical Viscoelastic Bio-Ink Particles. Advanced Functional Materials. 2022;32(6):2109004.

82. Zhu J, Cai LH. All-aqueous printing of viscoelastic droplets in yield-stress fluids. Acta Biomaterialia. 2023 Jul 15;165:60–71.

83. Zhu J, He Y, Wang Y, Cai LH. Voxelated bioprinting of modular double-network bio-ink droplets. Nat Commun. 2024 Jul 13;15(1):5902.

84. Yang Y, Fathi P, Holland G, Pan D, Wang NS, Esch MB. Pumpless microfluidic devices for generating healthy and diseased endothelia. Lab Chip. 2019 Sep 27;19(19):3212–9.

